# Eating breakfast and avoiding the evening snack sustains lipid oxidation

**DOI:** 10.1101/2020.01.29.923417

**Authors:** Kevin Parsons Kelly, Owen P. McGuinness, Maciej Buchowski, Jacob J. Hughey, Heidi Chen, James Powers, Terry Page, Carl Hirschie Johnson

## Abstract

Circadian (daily) regulation of metabolic pathways implies that food may be metabolized differentially over the daily cycle. To test that hypothesis, we monitored the metabolism of older subjects in a whole-room respiratory chamber over two separate 56-h sessions in a random crossover design. In one session, one of the three daily meals was presented as breakfast whereas in the other session, a nutritionally equivalent meal was presented as a late-evening snack. The duration of the overnight fast was the same for both sessions. Whereas the two sessions did not differ in overall energy expenditure, the respiratory exchange ratio (RER) was different during sleep between the two sessions. Unexpectedly, this difference in RER due to daily meal timing was not due to daily differences in physical activity, sleep disruption, or core body temperature. Rather, we found that the daily timing of nutrient availability coupled with daily/circadian control of metabolism drives a switch in substrate preference such that the late-evening snack session resulted in significantly lower lipid oxidation compared to the breakfast session. Therefore, the timing of meals during the day/night cycle affects how ingested food is oxidized or stored in humans with important implications for optimal eating habits.

## INTRODUCTION

Developed countries are experiencing an epidemic of obesity that leads to many serious health problems, foremost among which are increasing rates of Type 2 diabetes, metabolic syndrome, cardiovascular disease, and cancer. While weight gain and obesity are primarily determined by diet and exercise, there is tremendous interest in the possibility that the daily timing of eating might have a significant impact upon weight management [1–3]. Many physiological processes display day/night rhythms, including feeding behavior, lipid and carbohydrate metabolism, body temperature, and sleep. These daily oscillations are controlled by the circadian clock, which is composed of an autoregulatory biochemical mechanism that is expressed in tissues throughout the body and is coordinated by a master pacemaker located in the suprachiasmatic nuclei of the brain (aka the SCN [1,4]). The circadian system globally controls gene expression patterns so that metabolic pathways are differentially regulated over the day, including switching between carbohydrate and lipid catabolism [1,3,5–9]. Therefore, ingestion of the same food at different times of day could lead to differential metabolic outcomes, e.g., lipid oxidation vs. accumulation; however, whether this is true or not is unclear.

Non-optimal phasing of the endogenous circadian system with the environmental day/night cycle has adverse health consequences. Shiftworkers are a particularly cogent example, for their work schedule disrupts the optimal relationship between the internal biological clock and the environmental daily cycle, and this disruption leads to well-documented health decrements [6,9–13]. A contributing culprit that is often implicated in shiftwork’s temporal disruption is the disturbance of eating patterns and preferences. In non-human mammals, a persuasive literature demonstrates that manipulating the timing of feeding relative to biological clock phase effectively controls obesity [1,14,15]. In particular, mice fed a high-fat diet on a restricted schedule maintain a healthy weight when fed only during their active phase, but become obese if the high-fat diet is present during the inactive phase, even though the long-term caloric intake and locomotor activity levels are comparable between day- vs. night-fed mice [14,15].

Can the timing of eating relative to our circadian cycle of metabolism and sleep also help to regulate lipid metabolism and body weight in humans? Eating late in the day is correlated with weight gain [16], and there is an oft-discussed debate whether skipping breakfast versus dinner reaps weight-control benefits [3,17]. While many factors can influence the timing of eating in everyday life [2,3,18,19], we decided to take an experimental approach to test the metabolic consequences of a straightforward exchange of equivalent nutritional intake between early morning (8 am) and bedtime (10 pm). While it is not feasible to do the 12-h reversal of feeding time that was tested with mice [14] because it would disrupt the consolidated sleep episode of humans, we focused upon a 4.5 h shift of feeding where human subjects ate either a ~700 kcal breakfast or an equivalent ~700 kcal late-evening snack. Not only are these two feeding schedules experimentally tractable for a human study, they are also commonly practiced by humans in everyday life (i.e., “skipping breakfast” and/or “late-evening snacking"). For each feeding schedule, we monitored the metabolism of our subjects for a 56-h stint in a whole-room calorimetry chamber to continuously measure their metabolic rate, respiratory exchange ratio (RER), carbohydrate oxidation, and lipid oxidation.

Previous studies of human metabolism for shorter monitoring periods (~24 h) suggested that overall 24-h energy expenditure was not significantly affected by either breakfast skipping or a late dinner [20,21]); however, those prior studies were performed on healthy Asian young adults of optimal BMI (18.5-25 kg/m^2^) for only ~24 h, which is inadequate to study a phenomenon based on circadian rhythmicity. Moreover, while differences in blood glucose levels were reported in those studies between breakfast-skipping or late-dinner sessions, lipid oxidation was either not affected (breakfast-skipping [21]) or counter-intuitively enhanced (late-dinner [20]). In this investigation, we monitored older Caucasian adults (aged 50 or above) of varying BMI because we reasoned they are more representative of populations at-risk for metabolic disorders in many developed countries than are young and healthy adults. Each subject underwent two 56-h (2.5 d) sessions in a whole-room respiratory chamber, and with a randomized crossover design, we compared the energy expenditure (metabolic rate) and RER of each subject when given a scheduled breakfast, lunch, and dinner (Breakfast Session) versus when they were given a lunch, dinner, and a late-evening snack (Snack Session).

While overall 24-h energy expenditure was similar in this group of older subjects, RER was significantly different between the two sessions. We anticipated that daily differences in physical activity, sleep disruption, or core body temperature might lead to differential metabolism as reflected in the RER. Unexpectedly, however, our data demonstrated that even though the total daily energy and nutrient intake was equivalent between the sessions, switching the daily timing of a nutritionally equivalent 700-kcal meal from a “breakfast” to a “late-evening snack” had a significant impact upon carbohydrate and lipid metabolism such that nocturnal carbohydrate oxidation was favored at the expense of lipid oxidation when subjects ate the 700-kcal meal as a late-evening snack. Therefore, the daily cycle of metabolism and nutrient availability switches substrate preference so that the cumulative net lipid oxidation is altered by the timing of meals.

## RESULTS

We studied the metabolism of human subjects by indirect calorimetry under continuous monitoring in Vanderbilt University’s Human Metabolic Chamber. During each visit, the minute-by minute oxygen consumption (VO_2_), carbon dioxide production (VCO_2_), actigraphy, and core body temperature (CBT) of the subjects were continuously measured, with subsequent calculation of RER (VCO_2_/VO_2_), metabolic rate, carbohydrate oxidation, and lipid oxidation. The subjects slept and ate in the metabolic chamber, and were allowed only two brief (20 min) episodes per day outside the chamber: once about 10:00 am to take a quick shower, and once about 3:00 pm to take a brief non-strenuous walk. Each subject was monitored for two full-duration 56-h experiments that compared differences in the timing of their meals. In the “Breakfast Session” (Fig. 1A), the subjects had breakfast, lunch, and dinner, with a ~13.75 hour fast from 6:15 pm (end of dinner) to 8:00 am (breakfast). In the “Snack Session,” subjects only had a cup of tea or coffee (without sugar or creamer) at breakfast time, and their first meal was lunch (Fig. 1A). Then, a snack was served at 10:00 pm just before sleep (lights-off), and the subjects fasted ~14 h until lunch (10:30 pm to 12:30 pm). The breakfasts and the snacks had equivalent nutritional and caloric values of ~700 kcal; the breakfast on Day 2 was identical to the snack on Day 2, and the breakfast on Day 3 was identical to the snack on Day 3 (Supplemental Table S1A). Therefore, the meals served to subjects during the Breakfast Session had equivalent energy and nutrient content as in the Snack Session over the 24-h day (see Supplementary Table 1 for detailed nutritional information). All subjects completed both sessions in this cross-over experiment, which allowed pairwise comparison of their data. Our study is distinguished from the earlier metabolic chamber studies of meal timing in humans [20,21] by the cross-over design of our protocol and the fact that we studied older subjects of various BMI (51-63 years old, BMI 22.2 - 33.4, see Table 1) who may be less resilient to metabolic perturbations for energy expenditure than are younger subjects with BMIs of 20-25 [22,23]. Moreover, our Breakfast vs. Snack Sessions had essentially the same duration of daily fasting (13.75-14 h) to avoid the confounding factor of differential fasting durations found in other studies [e.g., ref. 19].

**Figure 1.**
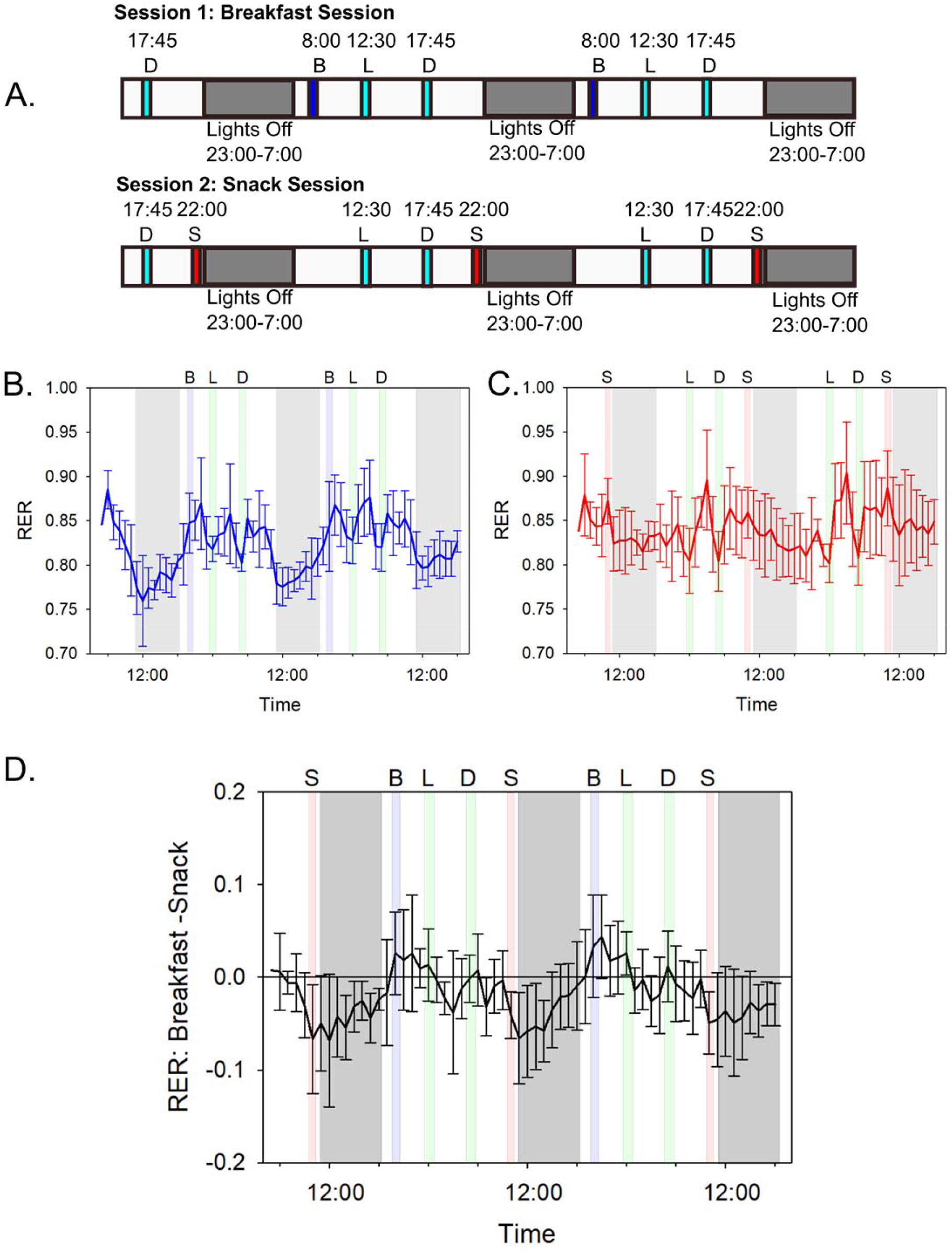
Chamber schedule and the impact of meal timing on subjects’ respiratory exchange ratios (RERs). See also Supplementary Table 4A. **A**) Protocol for “Breakfast” Session vs. “Snack” Session. Subjects experienced two separate 56-h continuous sessions with constant metabolic monitoring by indirect calorimetry, each session lasting 56 hours. The Breakfast session included a breakfast (B), lunch (L), and dinner (D) while the Snack session contained a lunch, dinner, and a late-evening snack (S). The late-evening snacks were of equivalent caloric and nutritional value to the breakfast meals (~700 kcal, see Supplementary Table 1A for details). Note from Supplementary Figure S1 that the daily phasing of sleep for the subjects prior to entry into the metabolic chamber was the same as the “lights-off” interval during the 56-h time course, so the subjects did not experience a phase shift of their daily cycle when they entered the experimental conditions. **B**) Breakfast Session: blue line indicates the average hourly RER over the entire 56-h time course among all six subjects when a breakfast, lunch, and dinner were presented. Error bars are the standard deviation. Letters indicate time and type of meals and gray shaded areas indicate the lights off periods. Green shaded areas indicate meals that were given at the same time in both Breakfast and Snack Sessions (lunch and dinner). Blue shaded areas indicate when breakfast was given, and gray shading indicates the lights off period. See Supplementary Figure S2 for data of all subjects individually. **C**) Snack Session: the red line indicates the average hourly RER over the entire 56-h time course among all subjects when a lunch, dinner, and a late-evening snack were presented. Red shaded areas indicate when late-evening snacks were given. Breakfasts and late-evening snacks contained the same amount of calories and the same lipid, carbohydrate, and protein content (Supplemental Table 1A/B). Error bars are the standard deviation (n=6). **D**) Average difference in RER over the entire 56-h time course for the Breakfast Session subtracted from the Snack Session. Deviation from zero (horizontal black line) indicates where differences in RER occurred between subjects. Error bars indicate standard deviation in the differences. In Panels B, C, & D, times of meals are indicated by letters (B = breakfast, L = lunch, D = dinner, S = late-evening snack), gray areas are lights-off (sleep) intervals. Meal times are shaded as in panels A-C; breakfasts and snacks occurred only in their respective sessions. All RER data were collected minute by minute, and in this figure the minute by minute data were binned and averaged for all 60 values within an hour. Abscissa are clock time.

**Table 1.**
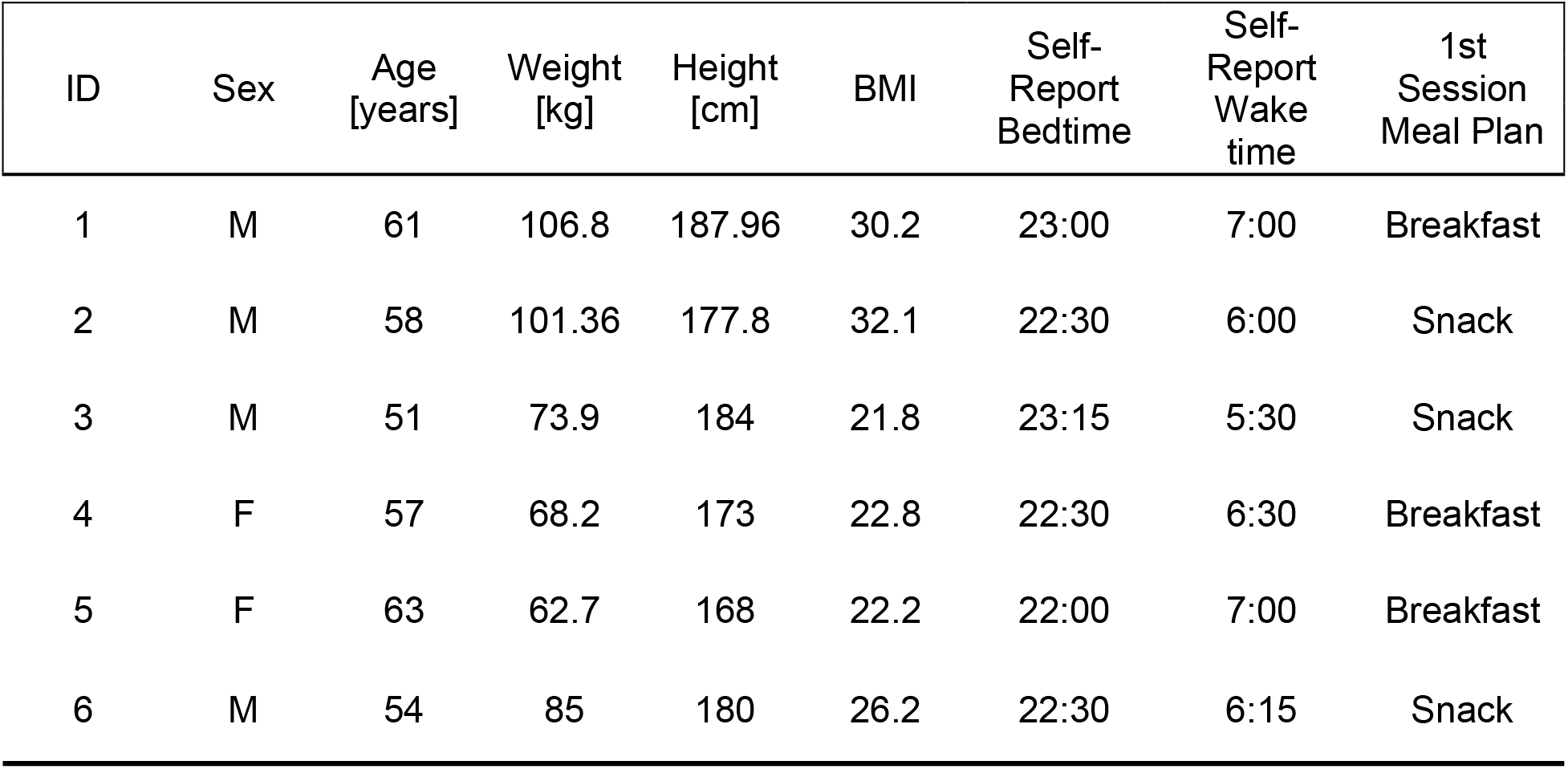
Subjects Involved in this study

| ID | Sex | Age<br>[years] | Weight<br>[kg] | Height<br>[cm] | BMI | Self-Report<br>Bedtime | Self-Report<br>Wake<br>time | 1st<br>Session<br>Meal Plan |
| --- | --- | --- | --- | --- | --- | --- | --- | --- |
| 1 | M | 61 | 106.8 | 187.96 | 30.2 | 23:00 | 7:00 | Breakfast |
| 2 | M | 58 | 101.36 | 177.8 | 32.1 | 22:30 | 6:00 | Snack |
| 3 | M | 51 | 73.9 | 184 | 21.8 | 23:15 | 5:30 | Snack |
| 4 | F | 57 | 68.2 | 173 | 22.8 | 22:30 | 6:30 | Breakfast |
| 5 | F | 63 | 62.7 | 168 | 22.2 | 22:00 | 7:00 | Breakfast |
| 6 | M | 54 | 85 | 180 | 26.2 | 22:30 | 6:15 | Snack |

As illustrated in Fig. 1B, subjects on the Breakfast Session that included breakfast and a fast throughout the time interval from dinner to the following breakfast (6:30 pm to 8:00 am), exhibited a strong daily rhythm of RER (aka Respiratory Quotient, calculated as VCO_2_ / VO_2_ [24,25]). RER values close to 0.7 indicate lipid oxidation while values of ~1.0 indicate almost exclusive carbohydrate oxidation. The average RER of this diet is similar to that of typical diets in the USA (~0.85); it includes a mixture of lipids, protein, and carbohydrates [24,25]. RER of subjects on the Breakfast Session was low throughout the lights-off interval (indicating primarily lipid catabolism during sleep) and high during the active daytime (indicating primarily carbohydrate and protein catabolism). Therefore, humans share with other mammals a daily rhythm of substrate metabolism as assessed by indirect calorimetry [26,27]. However, in the Snack Session, the metabolism of the same subjects was not as strongly rhythmic, and displayed a much lower amplitude rhythm of RER that did not drop into a largely lipid catabolic mode (Fig. 1C). When the difference between RER on the Breakfast vs. the Snack Sessions is calculated as a function of daily time, the most significant difference was noted during the inactive sleep phase where lipid oxidation predominates in the Breakfast Session while RER remains high in the Snack Session (Fig. 1D).

We initially predicted that the session-dependent RER patterns were due to differences between the sessions in physical activity, sleep disruption, core body temperature, or the phasing/amplitude of the circadian clock. However, none of these parameters were different between the sessions. Actigraphy confirmed that the subjects’ overall activity levels did not differ significantly between the two sessions (Fig. 2A/B, p = 0.538). Moreover, actigraphy can provide an assessment of restlessness during sleep [28] and by this criterion, the sleep quality was equivalent between the Breakfast vs. Snack Sessions (Fig. 2A/B). The daily rhythm of the core body temperature (CBT), that is frequently used as a marker of the central circadian clock in humans [9,29], did not show significant differences in the phasing or amplitude of their CBT rhythms between the two sessions (Fig. 2, p = 0.218). Moreover, the circadian rhythm of overall metabolic rate (MR, [8]) was not different in phase or amplitude between the two sessions (Fig. 3B, p = 0.11). Therefore, neither differences in core body temperature (Fig. 2) nor the thermic effect of food (TEF, see below) were responsible for the session-dependent RER differences. The phasing or amplitude of the daily rhythms of master clock markers (plasma melatonin and cortisol), insulin, or plasma triglycerides is also not responsible, as reported by Wehrens and co-workers who found no differences in those rhythms in a meal timing study using a very similar protocol to ours [30]. Finally, our subjects kept a regularly timed sleep/wake cycle prior to the metabolic chamber experiments so that their internal rhythm was in phase with the light/dark cycle during the 56-h experiment (Supplementary Figure 1, Table 1). These results suggest that the change in meal timing altered the RER rhythm (i.e., its amplitude) without changing overall activity, sleep quality, body temperature, or the phase relationship between circadian rhythms and the daily light/dark schedule.

**Figure 2.**
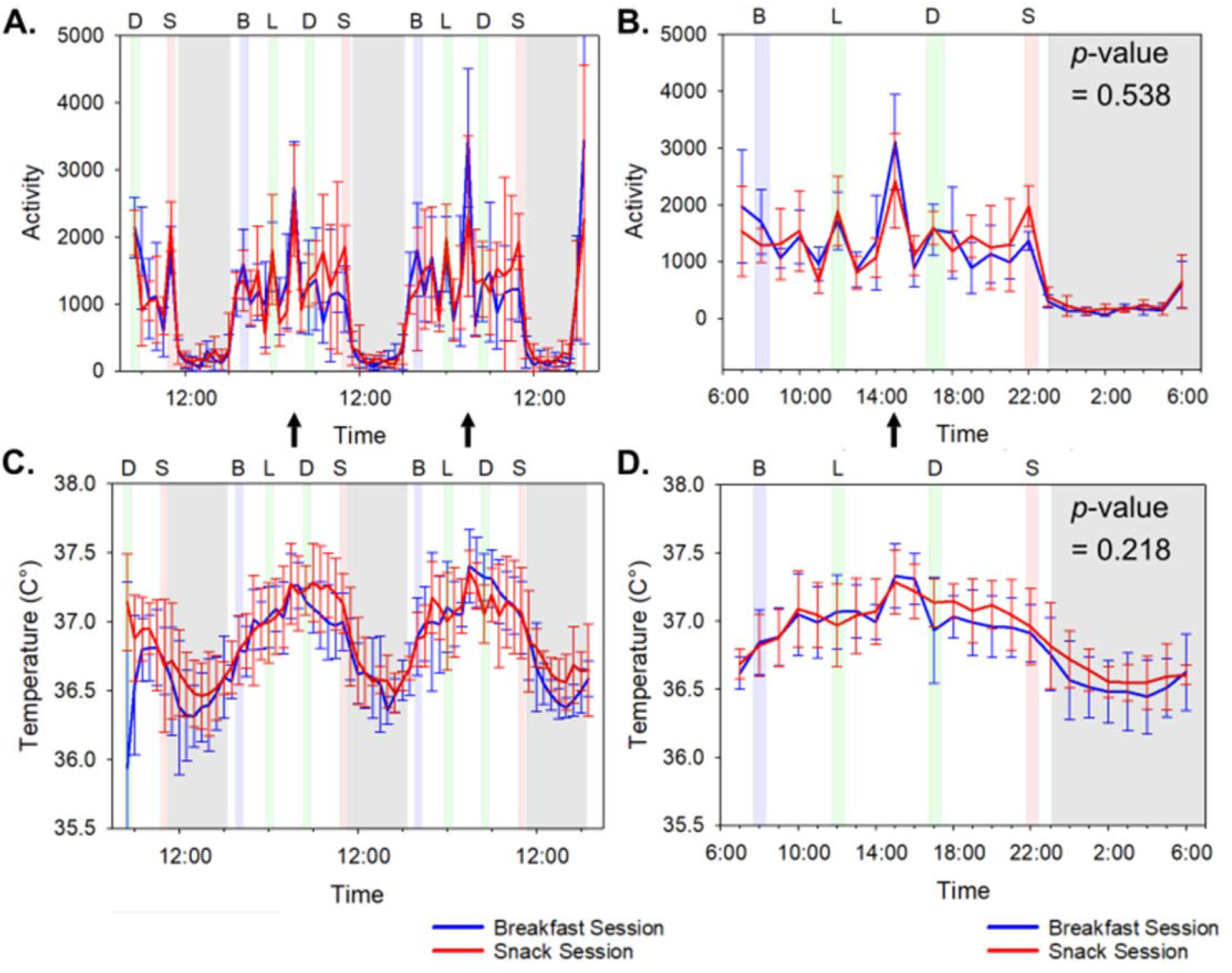
Activity and Core Body Temperature Patterns. See also Supplementary Table 4B and 4C. **A**) Average wrist locomotor activity (measured as vector of magnitude) of all subjects for the 56-h time course. The blue line indicates values during the subjects’ Breakfast sessions and the red line for the subjects’ Snack sessions. Black arrows indicate the afternoon break where subjects were allowed to exit the chamber for a 30 minute break, during which the subjects were allowed a non-strenuous walk. (This 30-min interval was excluded in other measurements as calorimetric readings were not being taken during this break.) **B**) Average wrist activity of all subjects plotted modulo-24 hours. Minute-by-minute activity data were averaged for all subjects into one hour bins and aligned by clock time. The arrow denotes the 30 min break referred as noted in panel A. The p-value of 0.538 refers to a pairwise comparison of the average (breakfast–snack) values over the full 56 h time course for wrist activity. See Supplementary Table 4B for the hour-by-hour statistical comparison of the breakfast versus the snack sessions. **C**) Average core body temperature (CBT) for all subjects over the 56-h time course. **D**) Average CBT of all subjects plotted modulo-24 hours. Minute-by-minute activity data were averaged for all subjects into one hour bins and aligned by clock time. The p-value of 0.218 refers to a pairwise comparison of the average (breakfast–snack values over the full 56-h time course for CBT. See Supplementary Table 4C for the hour-by-hour statistical comparison of the breakfast versus the snack sessions. **All panels:** The blue line indicates values during the subjects’ Breakfast sessions and the red line for the subjects’ Snack sessions. Shading indicates meals and lights-off as in Fig. 1B/C. Error bars indicate +/− standard deviation (n = 6).

The differences in the RER patterns between the two sessions manifest primarily during the time of late-evening snacking and for at least several hours into the sleep episode (hours 22 – 03 (Fig. 3E, Supplementary Table 4A). Apparently the late-evening snacking delays the clock-induced switching between primarily carbohydrate-catabolic mode (higher RER values) and primarily lipid-catabolic mode (lower RER values). Despite this change in the temporal pattern of RER, the values integrated over the entire 56-h time courses indicate differences slightly above the p = 0.05 level between the two sessions in terms of overall RER or total energy expenditure (Fig. 3C/F; p = 0.068 for RER and p = 0.11 for MR). Moreover, while there was a significant thermic effect of food (TEF) for MR and RER at each meal, our calculations based on the method of McHill and coworkers [11] indicated no differences in the TEF between the sessions for lunch and dinner–the two meals that were the same in both sessions (p = 0.432 for lunch and p = 0.855 for dinner). Moreover, the TEF for the breakfast as compared with the snack was also not different (p = 0.284). Therefore, differences in metabolic rates as assessed by TEF were not responsible for the substrate-switching preferences that are described below for breakfast-skipping versus late-evening snacking.

**Figure 3.**
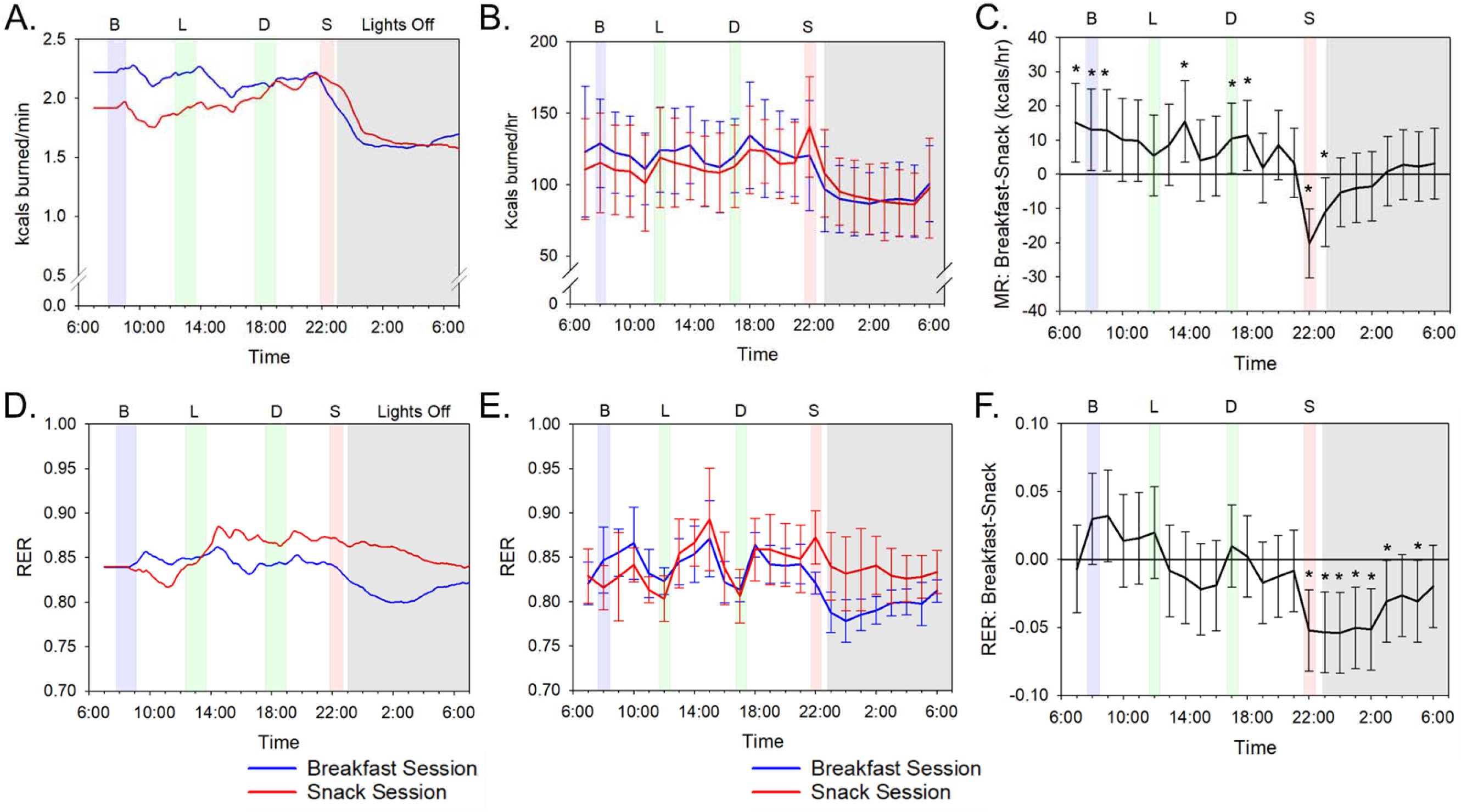
Metabolic Rates and RER Values. Refer to Supplementary Table 4A and 4D. **A**) Metabolic rate by indirect calorimetry for a representative participant (Subject #3). The data for Subject #3 are plotted as a moving average using 180 data points (= 3 h) after aligning all time points to clock time and integrated on a 24-h scale. **B**) Average metabolic rate for all subjects plotted modulo-24 hours. Data were averaged into 1-h bins with error bars indicating standard deviation. See Supplementary Figure S3 for data of all subjects’ metabolic rates individually plotted. **C**) Average hourly pairwise comparison of (breakfast–snack) metabolic rate values for all subjects. Error bars indicate 95% confidence intervals and values are based on a mixed model analysis. Asterisks indicate significant differences (p-value<0.05) between breakfast and snack values for the indicated one-hour bins. See Supplementary Table 4D for the hour-by-hour statistical comparison of the breakfast versus the snack sessions. **D**) RER (VCO_2_/VO_2_) by indirect calorimetry of a representative individual (subject #3). The data for Subject #3 are plotted as a moving average using 180 data points (= 3 h) after aligning all time points to clock time and integrated on a 24-h scale. **E**) Average RER for all subjects plotted modulo-24 hours. Data were averaged into 1-h bins with error bars indicating standard deviation. See Supplementary Figure S2 for data of all subjects’ RER individually plotted. **F**) Average hourly pairwise comparison of (breakfast–snack) RER values for all subjects. Error bars indicate 95% confidence intervals and values are based on a mixed model analysis. Asterisks indicate significant differences (p-value<0.05) between breakfast and snack values for the indicated one-hour bins. See Supplementary Table 4A for the hour-by-hour statistical comparison of the breakfast versus the snack sessions. **All panels:** The blue line indicates values during the subjects’ Breakfast sessions and the red line for the subjects’ Snack sessions. Shading indicates meals and lights-off as in Fig. 1B/C. Error bars indicate +/− standard deviation.

The conclusion that altered meal timing delays the sleep-onset switching between carbohydrate and lipid catabolic modes can be more easily visualized by converting the RER values into carbohydrate vs. lipid oxidation rates [25]. Carbohydrate oxidation during the Breakfast Session was high during the active day-phase with peaks just after each mealtime, but it dropped precipitously as the subjects entered their sleep episode after lights-out at 11:00 pm (Fig. 4A). The carbohydrate oxidation rate of subjects on the Breakfast Session who had not eaten since dinnertime began to fall before sleep onset and continued to be low through the first half of the nocturnal sleep episode (Supplementary Table 4E). On the other hand, in the Snack Session, the late-evening snack caused a peak carbohydrate catabolism just before going to bed, and while carbohydrate oxidation dropped thereafter, it remained higher throughout the sleep episode than when the same subjects were on the Breakfast Session (Fig. 4A). Overall, 24-h carbohydrate oxidation did not differ between sessions because the increased oxidation after breakfast in the Breakfast Session was offset by less carbohydrate oxidation in the early night of the Breakfast Session (Fig. 4C; Supplementary Table 4E). Therefore, carbohydrate oxidation was not significantly different between the sessions over the entire 56-h time course (p = 0.130).

**Figure 4.**
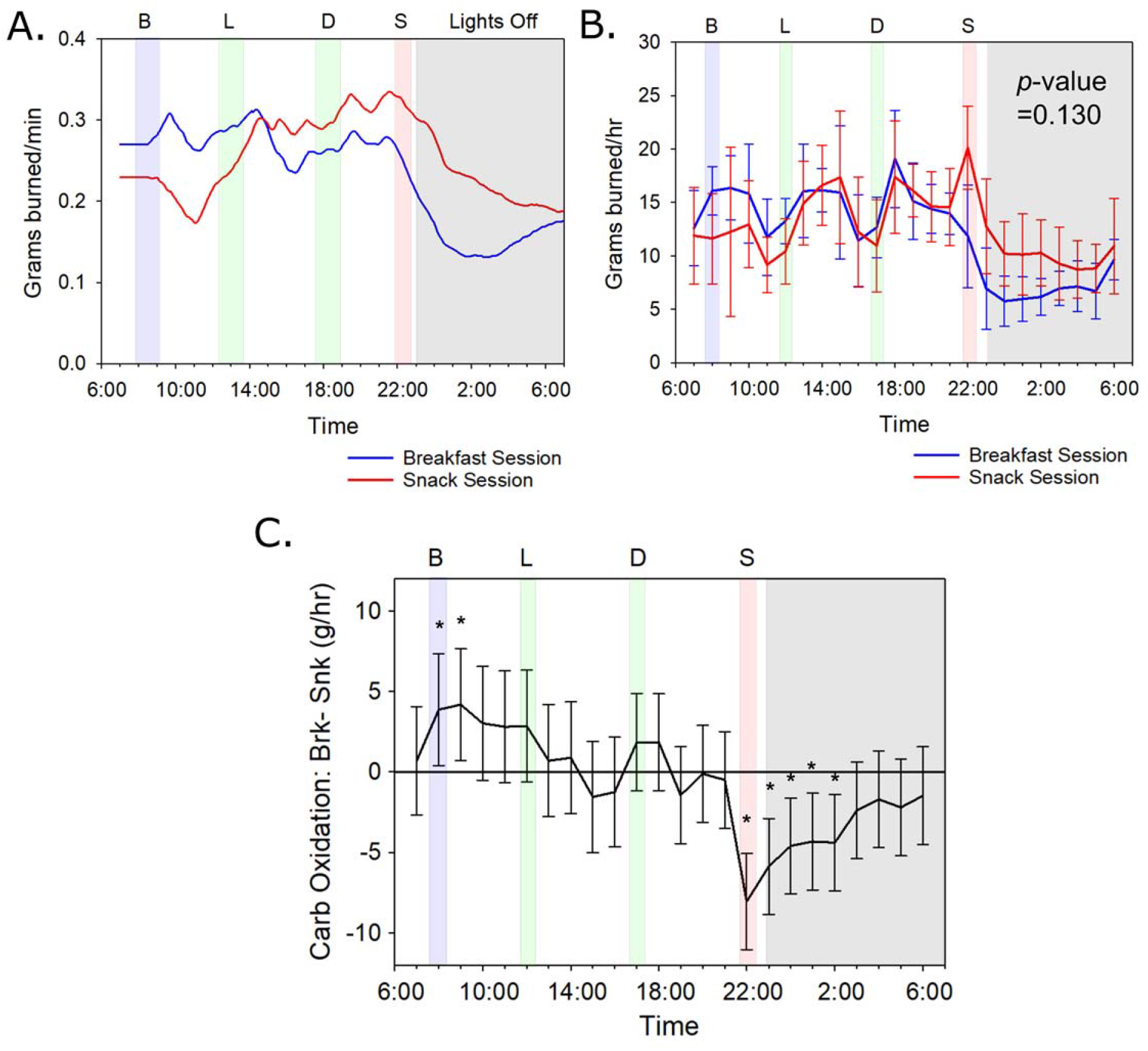
Carbohydate Oxidation Patterns Offset Between the Meal Timing Sessions; See also Supplementary Table 4E. **A**) Carbohydrate oxidation data of a representative subject (#3) calculated from indirect calorimetry measurements as described [24,25]. Data are plotted as a 3-h moving average (180-minute data points). **B**) Average of all subjects for daily carbohydrate oxidation calculated from indirect calorimetry measurements as described [24,25] and the 56-h time course data are plotted on a modulo-24 h scale. Averaged data for all subjects are organized in 1-h bins. The p-value of 0.130 refers to a pairwise comparison of the average (breakfast - snack) values over the full 56-h time course for carbohydrate oxidation. See Supplementary Figure S4 for data of all subjects’ carbohydrate oxidation rates plotted individually. **C**) Average hourly pairwise comparison of (breakfast–snack) carbohydrate oxidation values for all subjects. Error bars indicate 95% confidence intervals and values are based on a mixed model analysis. Asterisks indicate significant differences (p-value<0.05) between breakfast and snack values for the indicated one-hour bins. See Supplementary Table 4E for the hour-by-hour statistical comparison of the breakfast versus the snack sessions. **All panels:** The blue line indicates values during the subjects’ Breakfast sessions and the red line for the subjects’ Snack sessions. Shading indicates meals and lights-off as in Fig. 1B/C. Error bars indicate +/− standard deviation.

On the other hand, lipid oxidation was different between the sessions (p = 0.028). Subjects on the Breakfast Session experienced a relatively constant rate of lipid oxidation throughout the 24-h cycle (Fig. 5B). Because the overall metabolic rate declined during the night (Fig. 3A), this means that carbohydrate catabolism was “switched off” and lipid oxidation was sustained metabolism during the nightly fast. However, on the Snack Session, the availability of carbohydrates that was enabled by the late-evening snack supported metabolism during the night by carbohydrate oxidation; since the nocturnal metabolic rate is lower than the diurnal metabolic rate and carbohydrate catabolism is maintaining nocturnal metabolism, the meal timing of the Snack Session inhibited lipid oxidation at night (Fig. 4B/C). On average, 15 more grams of lipid were burned over the 24-h cycle by subjects on the Breakfast Session as compared with the Snack Session (Fig. 4D; p = 0.028).

**Figure 5.**
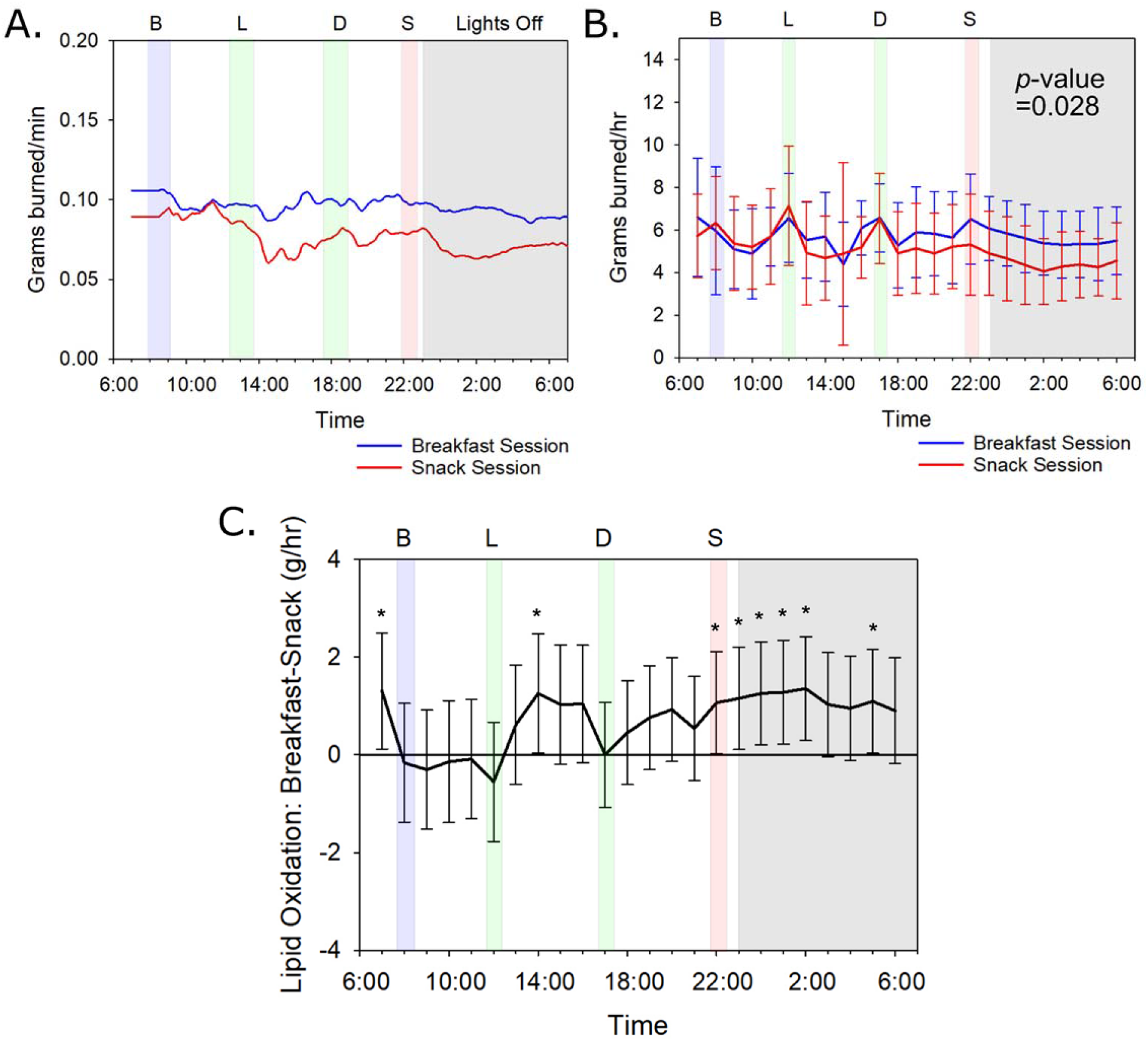
Meal Timing Alters Overall Lipid Oxidation; Refer to Refer to Supplementary Table 4F. **A**) Lipid oxidation data of a representative subject (#3) calculated from indirect calorimetry measurements as described [24,25]. Data are plotted as a 3-h moving average (180-minute data points). **B**) Average of all subjects for daily lipid oxidation calculated from indirect calorimetry measurements as described [24,25] and the 56-h time course data are plotted on a modulo-24 h scale. Averaged data for all subjects are organized in 1-h bins. The p-value of 0.028 refers to a pairwise comparison of the average (breakfast–snack) values over the full 56-h time course for lipid oxidation. See Supplementary Figure S5 for data of all subjects’ lipid oxidation rates plotted individually. **C**) Average hourly pairwise comparison of (breakfast–snack) lipid oxidation values for all subjects. Error bars indicate 95% confidence intervals and values are based on a mixed model analysis. Asterisks indicate significant differences (p-value<0.05) between breakfast and snack values for the indicated one-hour bins. See Supplementary Table 4F for the hour-by-hour statistical comparison of the breakfast versus the snack sessions. **All panels:** The blue line indicates values during the subjects’ Breakfast sessions and the red line for the subjects’ Snack sessions. Shading indicates meals and lights-off as in Fig. 1B/C. Error bars indicate +/− standard deviation.

Using a mixed model pairwise hour-by-hour analysis, we found significant differences in both carbohydrate and lipid oxidation in hours 22:00-02:00 of the snack/night interval (Supplementary Table 4E/F), with carbohydrates being utilized at higher rates for a longer time over this temporal window in the Snack Session than in the Breakfast Session (Fig. 4B/C). Conversely, the Breakfast Session showed significantly more lipids burned in the snack/night interval than did the Snack session (Fig. 5B/C). During the breakfast interval, we also found a significant difference in carbohydrate oxidation with more carbohydrates burned during the Breakfast Session than during the Snack Session at hours 08:00-09:00 (Supplementary Table 4E). Nevertheless, over the entire 24-h span there is not a net difference in carbohydrate oxidation between sessions because the oxidation difference in the breakfast window is offset by opposite oxidation rates in the snack/night window (Fig. 4C). However, the enhanced lipid oxidation in the snack/night window of subjects in the Breakfast Session is not offset by an opposite effect in another window (Fig. 5C). These results indicate that the time of meal placement can cause variation in the amount of lipids oxidized regardless of the nutritional or caloric content of the meal; changing the daily timing of a nutritionally equivalent meal of 700 kcal has a significant impact upon carbohydrate and lipid metabolism.

## DISCUSSION

The major finding of this study is that the timing of feeding over the day leads to significant differences in the metabolism of an equivalent 24-h nutritional intake. Daily timing of nutrient availability coupled with daily/circadian control of metabolism drives a switch in substrate preference such that the late-evening snack session resulted in significantly lower lipid oxidation compared to the breakfast session. When the subjects started bedrest after having just eaten the late-evening snack (Snack Session), they catabolized less lipid during their sleep episode than they did when they fasted from dinner to breakfast (Breakfast Session). This significant (p = 0.028, Fig. 5B) effect was measurable over only three sleep episodes in our experiments so that an average of 15 fewer grams of lipid were burned over the 24-h cycle by subjects on the Snack Session. The impact of a regular late-evening snack persisting over a longer time would progressively lead to substantially lower lipid oxidation (and therefore, more lipid accumulation) as compared with fasting during this interval of the day. As schematized in Figure 6, the daily patterns of substrate oxidation (Fig. 6A,6B) are roughly following the daily eating patterns (Fig. 6C). However, a late-evening snack likely sustains liver glycogen stores (carbohydrate oxidation, Fig. 6A) so that metabolism does not transition as rapidly or as fully into lipid oxidation during the nocturnal fast (Fig. 6B).

**Figure 6.**
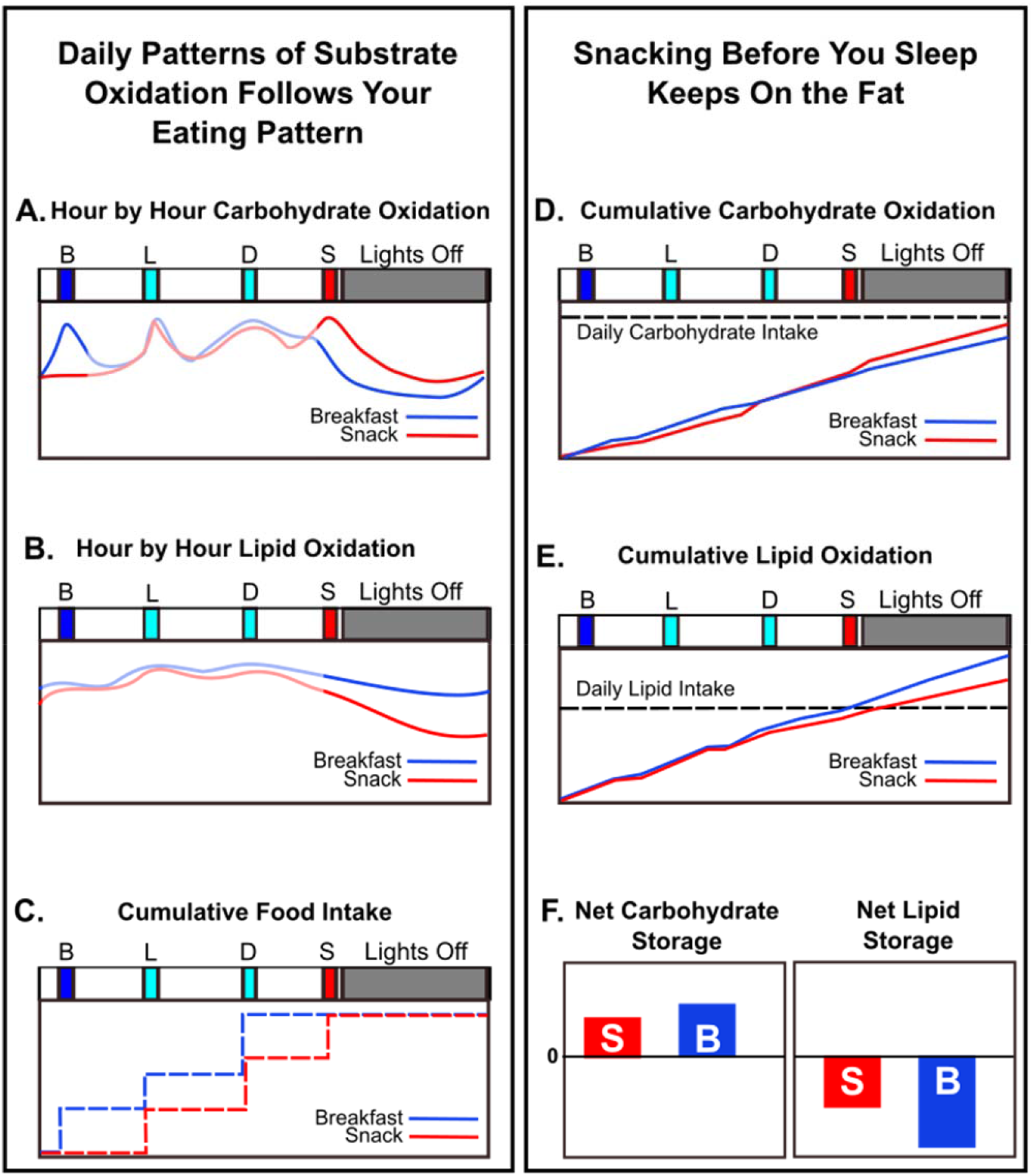
Schematic: late-evening snacking interacts with the circadian rhythm of metabolism to inhibit lipid oxidation. **A & B)** Hour by hour oxidation rates for carbohydrates (panel **A**) and lipids (panel **B**) in the two sessions. These curves are smoothed versions of the experimental data in Figures 4 & 5. **C)** Cumulative food intake on the Breakfast vs. Snack sessions. **D & E)** Cumulative oxidation rates over the 24-h cycle derived from the curves in panels **A** and **B**, and the experimental data of Figures 4& 5. Panel **D** shows cumulative carbohydrate oxidation, while panel **E** show cumulative lipid oxidation. The horizontal dashed lines indicate the daily total intake of carbohydrates (**D**) and lipids (**E**) for comparison with the cumulative respective oxidations. **F)** Approximate net relative daily storage of carbohydrates and lipids inferred from the data of Figures 4 & 5 and the analyses depicted in the other panels of this figure. Positive values indicate the extent of substrate accumulation/storage, negative values indicate the extent of substrate oxidation (“burning”).

Our interpretation of these data is based on the circadian clock orchestrating a switch between primarily carbohydrate oxidation to primarily lipid oxidation between the last meal of the day and the onset of circadian-timed sleep [1,3,6,7]. Instead of fasting between dinnertime and breakfast, if a person eats during the late evening, carbohydrates will be preferentially metabolized as sleep initiates, delaying the timing of the switch to primarily lipid oxidation. Over the 24-h cycle, cumulative carbohydrate oxidation as compared with total carbohydrate intake was not dramatically different between the two sessions (Fig. 6D), so the net 24-h carbohydrate storage is similar (Fig. 6F). On the other hand, the cumulative 24-h lipid oxidation rate as compared with total lipid intake is substantially less when late-evening snacking detains the transition to lipid catabolism (Fig. 6E), thereby lessening the mobilization of lipid stores (i.e., the extent of lipids being oxidized, Fig. 6F). There is a clear trade-off between lipid and carbohydrate oxidation during the night; the Breakfast Session clearly favors lipid oxidation at the expense of carbohydrate oxidation (Figs. 5 and 6).

There was a small but significant increase in carbohydrate oxidation in the morning after eating breakfast (Figs. 1D & 5A, hours 08:00-09:00 in Supplementary Table 4E) but not on lipid oxidation (Fig. 5B, hours 08:00-09:00 in Supplementary Table 4F). However, the effect of eating vs. skipping breakfast is not as significant as the impact of eating after dinner on both carbohydrate and lipid catabolism during the sleep episode (Fig. 5). In this study, there were no obvious differences among subjects based on BMI or gender (Table 1, Supplemental Figs. S2, S3, S4, S5). Unlike the conclusions of a previous investigation comparing morning versus evening carbohydrate-rich meals [31], in our investigation the different phasing of the meals between the two sessions did not change the phasing of the daily metabolic pattern. The phasing of sleep during the 56-h time courses matched that of the subjects’ sleep patterns for the prior week (compare Table 1 with Supplemental Fig. S1), and the phasing of the daily rhythms of activity, metabolic rate, and CBT were in phase between the two sessions (Figs. 2 and 3B). Therefore, in our protocol, the wake/sleep cycle appears to be locked in the same phase relationship to the lights-on/lights-off cycle in both sessions, and the altered meal timing of the Snack Session has delayed the metabolic switching between primarily carbo-catabolism mode and primarily lipid-catabolism modes in relationship to either the circadian system and/or the timing of sleep (Fig. 6).

Consistent with the findings of other investigations of altered meal timing, breakfast skipping, time-restricted feeding, etc. [19,20,21], we found no significant differences in total energy expenditure between sessions (Fig. 3C). Nevertheless, metabolism was significantly affected. In particular, the average daily RER maintained a higher value in the Snack Session (Fig. 3E/F), which can be attributed to a delayed entry into primarily lipid oxidation mode (Figs. 3E, 5, 6). The end result of the reduced lipid oxidation will be enhanced lipid storage, which over time will lead to increased adiposity. Therefore, in older adults who are potentially at-risk for metabolic disorders, avoiding snacking after the evening meal can sustain lipid oxidation and potentially improve metabolic outcomes.

## Materials and Methods

### Subjects

Six Subjects (4 Males and 2 Females) were first recruited through flyers and the Vanderbilt Kennedy Center. Subjects were between 51-63 (average age was 57) with Body Mass Indexes (BMIs) between 22.2-33.4 (Table 1). Applicants had to be 50 years of age or older, have no serious health complications or medications that could impact metabolism. Female subjects were not required to be post-menopausal for inclusion in this study, but because of the age requirement, all females recruited to the study were post-menopausal. Subjects had no prior shiftwork experience. The study protocol was approved by the Institutional Review Board of Vanderbilt University’s Human Research Protections Program (Approval number: 140536) and registered at ClinicalTrials.gov (identifier: NCT04144426). Prior to the study, each subject signed an informed and written consent.

Subjects were requested to monitor their feeding and sleeping habits for the one week prior to each metabolic chamber visit with a log that was provided. Subjects were asked to maintain their regular sleep and feeding schedule for one week prior to the chamber visits. Serendipitiously, all of the subjects maintained a typical sleeping and eating schedule that was approximately in phase with the meal and sleep (lights-off) schedule of the chamber visit (Table 1). Representative examples of the subjects’ sleep schedule prior to chamber visits appear in Supplementary Figure S1. After the one-week period, subjects were admitted to the Center for Clinical Research at Vanderbilt University after a health assessment by a physician. Metabolism of the subjects was monitored in the Human Metabolic Chamber at Vanderbilt University, which is a whole-room calorimeter with CO_2_ and O_2_ detectors to monitor the rate of VO_2_ and VCO_2_ (see Supplementary Figure S6). The room had a set flow rate of O_2_ and CO_2_ that allowed the energy expenditure of each subject to be measured through indirect calorimetry. Subjects were maintained on an enforced daily light/dark schedule where lights-on occurred at 7:00 am and lights-off at 11:00 pm. The subjects ate and slept in the metabolic chamber, and were allowed only two brief 20 min episodes per day outside the chamber: once about 10:00 am to take a quick shower, and once about 3:00 pm to take a brief non-strenuous walk. While in the chamber, the subjects were instructed to do sedentary activities like reading, internet, watching TV, etc.

During both visits, the Vitalsense^®^ Intergrated Monitoring Physiological System was used to monitor core body temperature (CBT) over the course of the study. Subjects were given a Vitalsense telemetric core body temperature capsule that recorded the subject’s core body temperature and relayed information to a monitor attached to the subject’s waistband (or under the pillow during lights-off). Telemetric capsules were given every 24 hours to maintain consistent temperature readings independent of bowel movements when the sensor might be excreted. Meals and lights-off times were scheduled regularly as shown in Figure 1A. Subjects were admitted into the metabolic chamber for two and a half days starting at 5:30 pm and ending 7:00 am after the third night in the facility. Subjects were admitted for two separate sessions at the facility with a shifted meal schedule. For one session (the Breakfast Session), subjects were given a breakfast at 8:00 am, lunch at 12:30 pm, and dinner at 5:45 pm every day. For the other session (the Snack Session), subjects were given decaf coffee (without cream or sugar) at 8:00 am, lunch at 12:30 pm, dinner at 5:45 pm, and a late-evening snack at 10:00 pm. The breakfast on Day 2 was identical to the snack on Day 2, and the breakfast on Day 3 was identical to the snack on Day 3 (Supplemental Table S1A), and each were ~700 kcal. The menus of the meals are shown in Supplemental Table S1A. The order of the sessions (i.e., Breakfast Session first versus Snack Session first) was determined in a randomized design for each subject (Table 1), and 4-12 days elapsed between sessions, depending upon the subject. Subjects were asked to eat all the meal provided but any leftover food was weighed back and actual intake for each meal is shown in Supplemental Table 1B. The size of the meals was determined by nutritionist to account for calories burned for each individual (on average, a daily 2300 kcal diet). Calories were divided as follows: ~700 kcals for Breakfast/Snack, ~600 kcals for lunch, and ~1000 kcals for Dinner.

### Whole-room Respiratory Chamber

The room calorimeter at Vanderbilt University is an airtight room (17.9 m³) providing an environment for daily living whose accuracy has been documented (Supplementary Figure S6 [32]). The room has an entrance door, an air lock for passing food and other items, and an outside window. The room is equipped with a TV/media system, toilet, sink, desk, chair, and rollaway bed allowing overnight stays. The calorimeter is located in the Clinical Research Center at Vanderbilt University and an intercom connects the chamber to a nearby station where nurses are on duty 24 hours/7 days per week. Temperature, barometric pressure, and humidity of the room are controlled and monitored. Minute-by-minute energy expenditure (kcal/min) are calculated from measured rates of O_2_ consumption and CO_2_ production using Weir’s equation [33].

### Quantification and Statistical Analyses

To quantify the differences between the Breakfast and Snack sessions, we applied a linear mixed model to the full 56-h time-course for each of the following measurements: metabolic rate (MR), activity, carbohydrate oxidation (CO), lipid oxidation (LO), Respiratory Exchange Ratio (RER), and Core Body Temperature (CBT). Each measurement was averaged using hourly bins for each subject in each session. By using a mixed model, we were able to adjust for dependency of within-subject observations. The model included fixed effects for session (Breakfast vs. Snack), day (treated as a factor variable and defined as 3:00 pm on one day to 2:59 pm on the next day), hour (treated as a factor variable), an interaction between session and day, and an interaction between session and hour. If the p-value of an interaction was greater than 0.2, we removed that interaction from the model.

### Calculations

Daily carbohydrate oxidation and daily lipid oxidation was calculated from indirect calorimetry measurements as described [24,25]. Nitrogen excretion rate was based on the amount on protein provided to subjects as well as previous research that monitored 24-hour nitrogen using similar parameters [34]. Because protein content was not altered between sessions, we assumed that 24-h nitrogen excretion rate was equivalent for Breakfast vs. Snack Sessions.

## Acknowledgements

We dedicate this study to the memory of Dr. Martin Katahn, a pioneer of the Rotation Diet and the benefactor whose donation enabled the metabolic chamber at Vanderbilt University. The authors would like to thank Briana Wyzinski, Ian Dew, and Regina Tyree for their contributions to subject recruitment and respiratory chamber operation. We thank Cynthia Dossett, Holly Mason, and Dr. Heidi Silver for their help with nutrition information and meal development. We thank Dr. Heidi Chen for her statistical advice. We also thank the subject-participants in this study. This study was supported by a Vanderbilt Discovery Grant, the Vanderbilt Institute for Clinical and Translational Research (VICTR) award ID# VR9806, the Vanderbilt Diabetes Research and Training Center (through the Metabolic Physiology Shared Resource supported by P60-DK020593), and the National Institute of Neurological Disorders and Stroke (R01 NS104497).

## Author Contributions

Experimental design: CHJ, TP, OM, MB

Subject recruitment and health assessement: KK, JP Performance of the experiments: KK, MB

Data analysis: KK, CHJ, TLP, OM, MB, JH

Wrote and edited the manuscript: KK, CHJ, TLP, OM, MB, JH

