## Supplemental Figures 1-6 and Supplemental Tables 1-4 for "Eating breakfast and avoiding the evening snack sustains lipid oxidation"

**By Kelly et al.**

##### Legends for Supplemental Figures

**Supplemental Figure S1 (Related to Table 1 and Fig. 1). Subject's self-reported schedules prior to entry into experiment.** The subjects' self-reported bedtime and wake-up time for the week prior to entry into the metabolic chamber shows that the daily phasing of sleep was similar before and during the 56-h experimental time course. Black squares specify the time of bedtime and wake-up, with the horizontal lines indicating sleep episodes prior to entry into the metabolic chamber (blue horizontal lines) or during the 56-h experimental time course (red horizontal lines). Therefore, the subjects did not experience a phase shift of their daily cycle when they entered the experimental conditions in the metabolic chamber.

**Supplemental Figure S2 (Related to Figs. 1 and 3). Individual Daily Respiratory Exchange Ratio (RER) data.** Average RER ($VCO_2/VO_2$) values for Subjects 1-6 from their Breakfast session (blue) and Snack Session (red) averaged into 1 hour intervals. Error bars indicate standard deviation.

**Supplemental Figure S3 (Related to Fig. 3). Individual Daily Metabolic Rate (MR) data.** Hourly kcals burned for Subjects 1-6 from their Breakfast session (blue) and Snack Session (red) averaged into 1 hour intervals. Error bars indicate standard deviation.

**Supplemental Figure S4 (Related to Fig. 4). Individual Daily Carbohydrate Oxidation.** Hourly grams of carbohydrates burned for Subjects 1-6 from their Breakfast session (blue) and Snack Session (red) averaged into 1 hour intervals. Daily carbohydrate oxidation values calculated from indirect calorimetry measurements as described [24,25]. Error bars indicate standard deviation.

**Supplemental Figure S5 (Related to Fig. 5). Individual Daily Lipid Oxidation.** Hourly grams of lipids burned for Subjects 1-6 from their Breakfast session (blue) and Snack Session (red) averaged into 1 hour intervals. Daily lipid oxidation values calculated from indirect calorimetry measurements as described [24,25]. Error bars indicate standard deviation.

**Supplemental Figure S6 (Related to Methods). Configuration and photographs of the human whole-room calorimetry chamber at Vanderbilt University.**

**Supplemental Tables**

**Supplemental Table 1**

A. Representative Meals
B. Nutritional Information

**Supplementary Table 2.** Inclusion/Exclusion Criteria

**Supplemental Table 3.** Questionnaire for subject recruitment

**Supplemental Table 4. Hour by Hour Mixed Model Analyses for:**

**A. RER**
**B. Activity**
**C. Core Body Temperature**
**D. Metabolic Rate**
**E. Carbohydrate Oxidation**
**F. Lipid Oxidation**

### Supplemental Figures

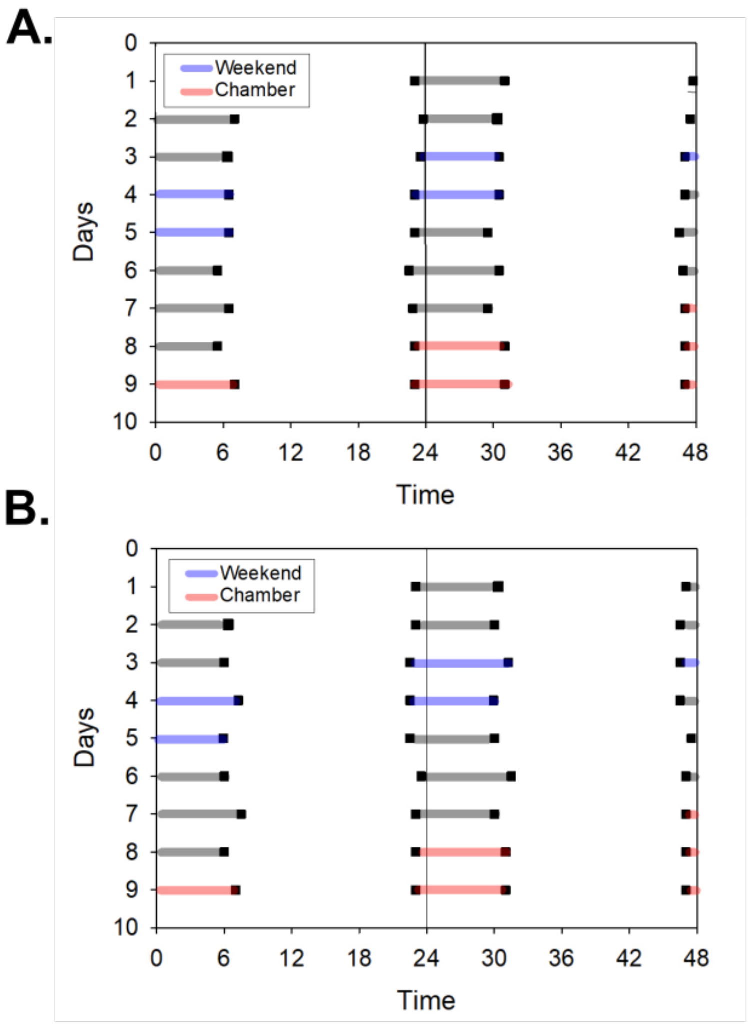

**Supplementary Figure 1. Representative Subject's self-reported sleep schedules prior to entry in the experiment for Breakfast (A) and Snack (B) Sessions.**

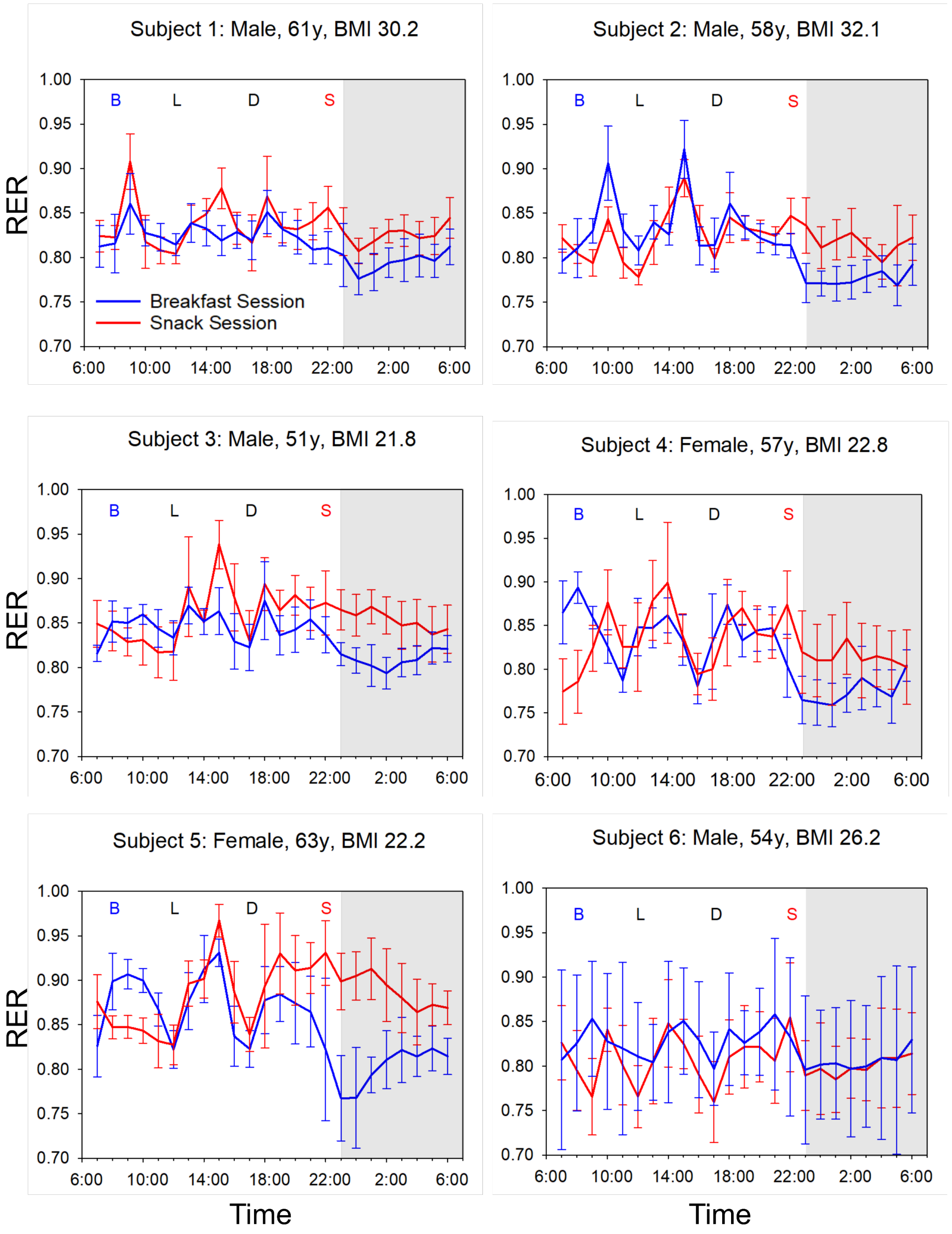

**Supplemental Figure 2. Daily Respiratory Exchange Ratio (RER) data for all subjects.**

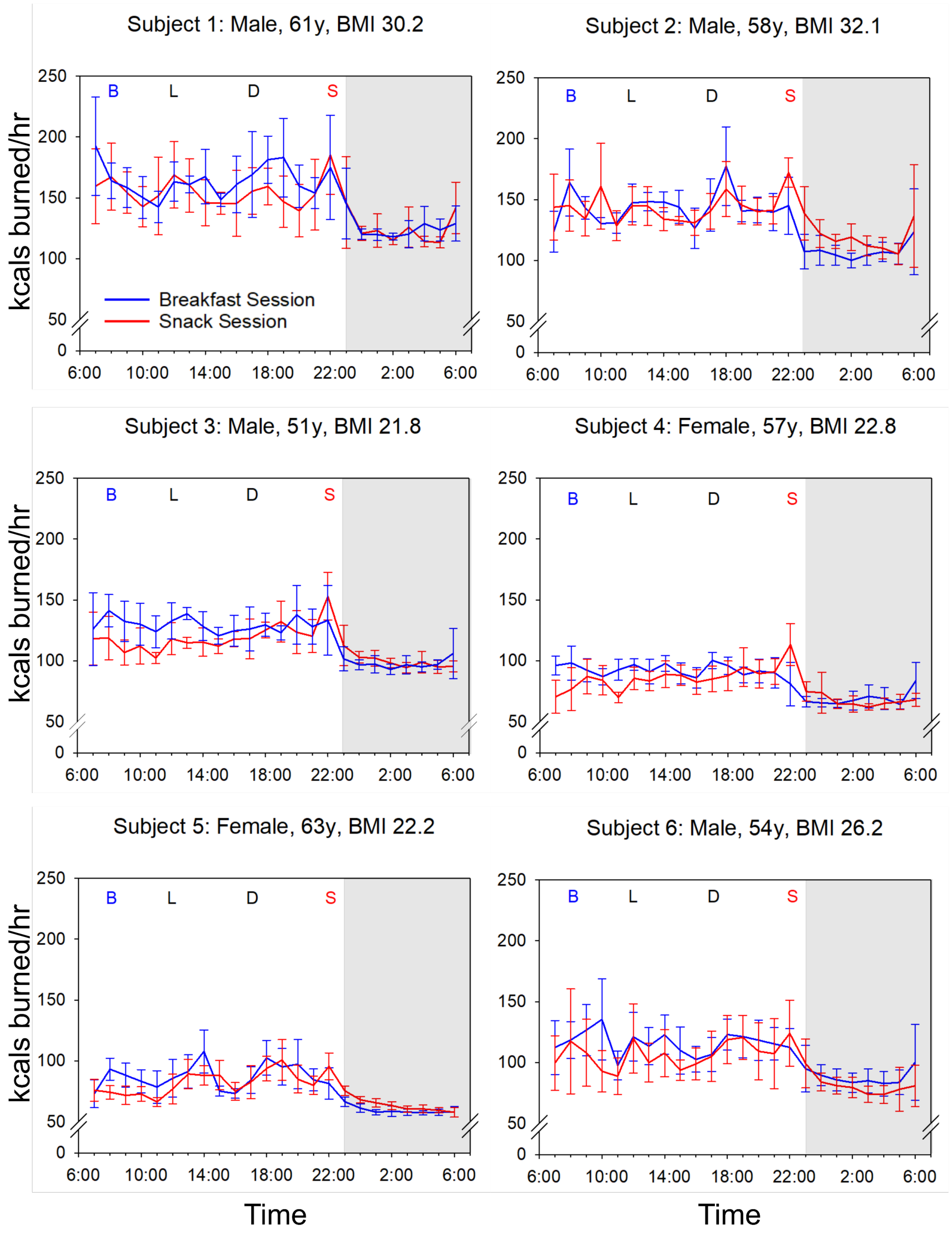

**Supplemental Figure 3. Daily Metabolic Rate data for all subjects.**

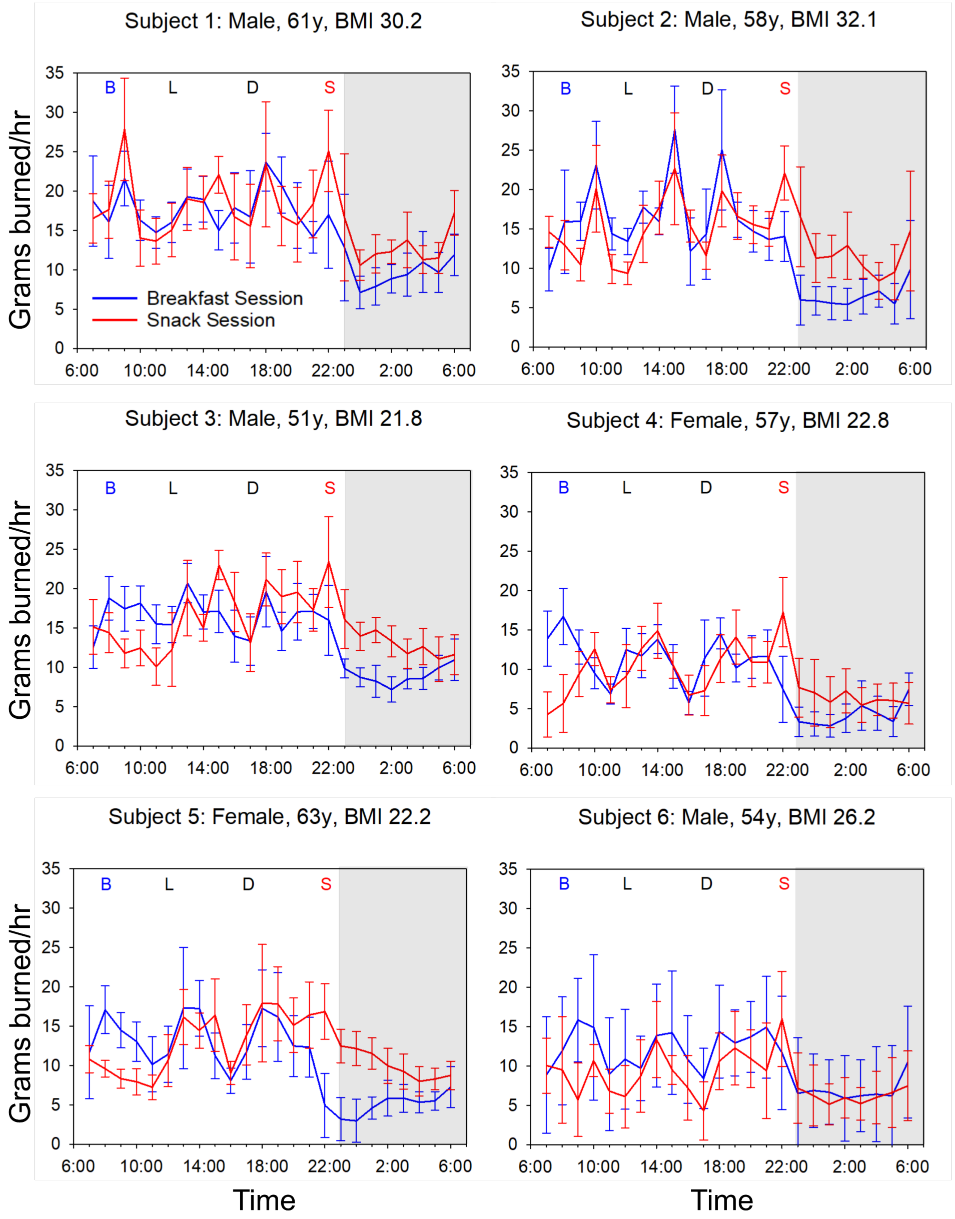

**Supplemental Figure 4. Daily Carbohydrate Oxidation data for all subjects.**

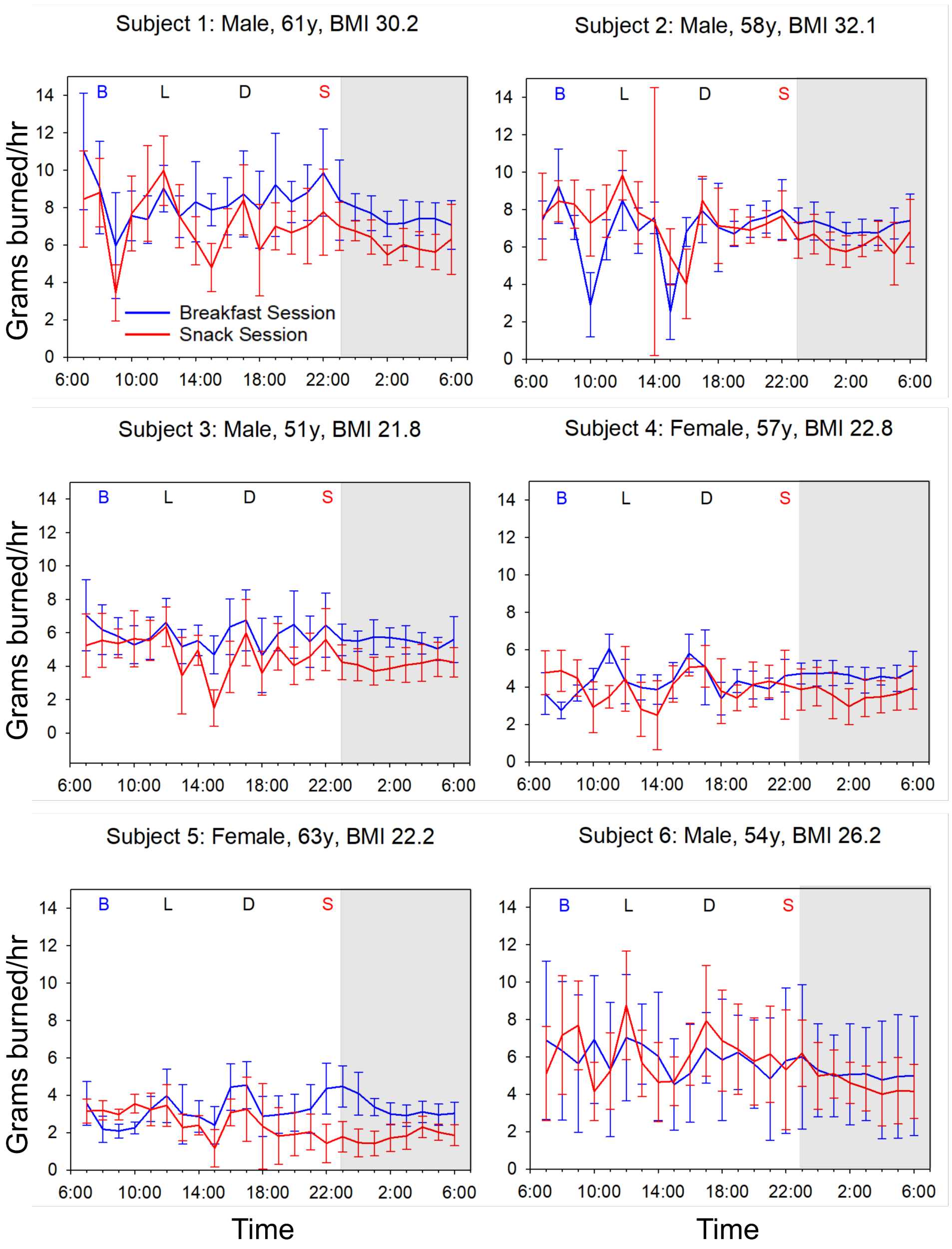

**Supplemental Figure 5. Daily Lipid Oxidation data for all subjects.**

#### Supplementary Figure S6

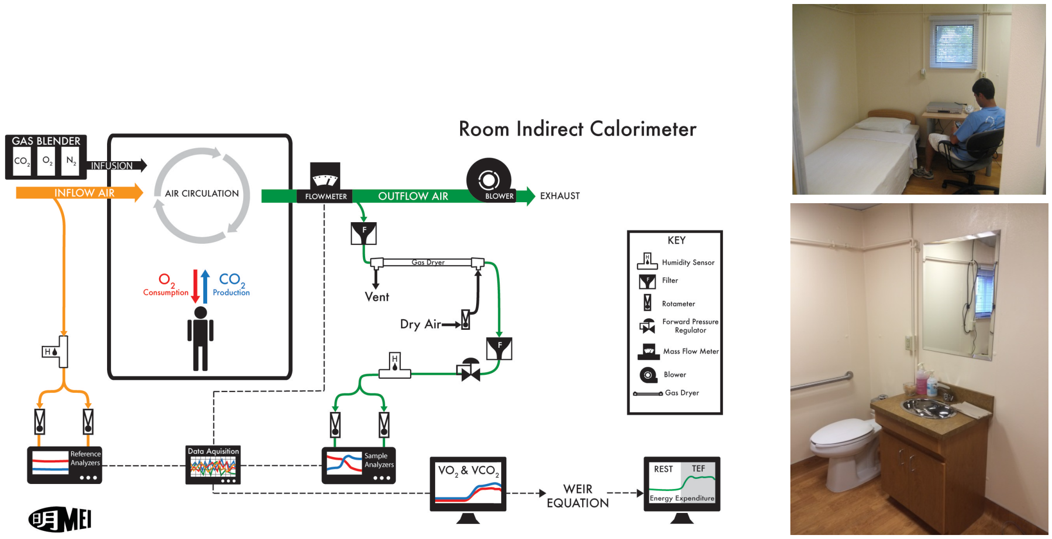

**Supplemental Figure 6. Human Metabolic Chamber at Vanderbilt University**

**Supplementary Table 1. Meals and Nutritional Information**

**A. Representative Meals**

| Breakfast Session | | | |
|---|---|---|---|
| **Admit Day 1** | **Day 2** | **Day 3** | **Day 4** |
| ***Breakfast*** | ***Breakfast 8AM*** | ***Breakfast 8AM*** | ***Breakfast 8AM*** |
| not applicable | English muffin / Margarine | Bagel / Cream cheese | Regular breakfast meal after |
| | Fresh pork patty | Boiled egg | study is completed. |
| | Honey nut cheerios | Apple | |
| | Mandarin orange | String cheese | |
| | Cranberry juice / 2% Milk | Orange juice | |
| | Decaf coffee or decaf tea | Decaf coffee or decaf tea | |
| ***Lunch*** | ***Lunch 12:30PM*** | ***Lunch 12:30PM*** | ***Lunch*** |
| not applicable | Turkey Sandwich | Hamburger | not applicable |
| | Salad / Ranch dressing | Steak house fries | |
| | Pears | Salad / Fat free Italian dressing | |
| | Potato chips | Peaches | |
| | Graham crackers | Grape juice | |
| ***Dinner 5:45PM*** | ***Dinner 5:45PM*** | ***Dinner 5:45PM*** | ***Dinner*** |
| Italian chicken | Beef Roast / Gravy | Roast turkey / Gravy | not applicable |
| Mashed potatoes | Scalloped potatoes | Brown rice | |
| Green beans / Margarine | Peas & carrots | Broccoli | |
| Grapes | Dinner roll / Margarine | Dinner roll / Butter | |

| | | |
|---|---|---|
| Snackwell cookies | Cantaloupe | Grapes |
| Sprite | Yogurt | Raspberry bar |
| | Peanut butter cookie | Apple juice |
| | Grape juice | |

| **Snack Session** | | | |
|---|---|---|---|
| **Admit Day 1** | **Day 2** | **Day 3** | **Day 4** |
| ***Breakfast*** | ***Breakfast 8AM*** | ***Breakfast 8AM*** | ***Breakfast 8AM*** |
| not applicable | Decaf coffee or decaf tea | Decaf coffee or decaf tea | Regular breakfast meal after |
| | | | study is completed. |
| ***Lunch*** | ***Lunch 12:30PM*** | ***Lunch 12:30PM*** | ***Lunch*** |
| not applicable | Turkey Sandwich | Hamburger | not applicable |
| | Salad / Ranch dressing | Steak house fries | |
| | Pears | Salad / Fat free Italian dressing | |
| | Potato chips | Peaches | |
| | Graham crackers | Grape juice | |
| ***Dinner 5:45PM*** | ***Dinner 5:45PM*** | ***Dinner 5:45PM*** | ***Dinner*** |
| Italian chicken | Beef Roast / Gravy | Roast turkey / Gravy | not applicable |
| Mashed potatoes | Scalloped potatoes | Brown rice | |
| Green beans / Margarine | Peas & carrots | Broccoli | |
| Grapes | Dinner roll / Margarine | Dinner roll / Butter | |

| | | | |
|---|---|---|---|
| Snackwell cookies | Cantaloupe | Grapes | |
| Sprite | Yogurt | Raspberry bar | |
| | Peanut butter cookie | Apple juice | |
| | Grape juice | | |
| ***Snack 11:00PM*** | ***Snack 11:00PM*** | ***Snack 11:00PM*** | ***Snack*** |
| Roast Beef Sandwich | English muffin / Margarine | Bagel / Cream cheese | not applicable |
| Orange | Fresh pork patty | Boiled egg | |
| Potato chips | Honey nut cheerios | Apple | |
| Graham crackers | Mandarin orange | String cheese | |
| Apple juice | Cranberry juice / 2% Milk | Orange juice | |

**B. Nutritional Summary**

**2500 kcal Diet**

| Meal Name | Avg. Total Grams | Avg. Energy (kcal) | Avg. Total Fat (g) | Avg. Total Carbohydrate (g) | Avg. Total Protein (g) |
|---|---|---|---|---|---|
| **Breakfast** | 824.1 +/- 107.5 | 750.0 +/- 22.6 | 24.9 +/- 1.7 | 103.6 +/- 4.8 | 28.6 +/- 2.7 |
| **Lunch** | 624.1 +/- 121.3 | 691.9 +/- 29.6 | 20.6 +/- 29.6 | 105.3 +/- 7.5 | 26.5 +/- 7.7 |
| **Dinner** | 883.9 +/- 116.0 | 937.6 +/- 53.2 | 26.5 +/- 4.3 | 139.0 +/- 12.3 | 43.8 +/- 6.0 |
| **Snack** | 608.4 +/- 122.0 | 746.2 +/- 65.9 | 25.1 +/- 2.5 | 103.9 +/- 9.9 | 28.1 +/- 3.2 |
| **Beverage Only** | 236.3 +/- 7.8 | 3.1 +/- 0.1 | 0 +/- 0 | 0.7 +/- 0 | 0 +/- 0 |

**2000 kcal Diet**

| Meal Name | Avg. Total Grams | Avg. Energy (kcal) | Avg. Total Fat (g) | Avg. Total Carbohydrate (g) | Avg. Total Protein (g) |
|---|---|---|---|---|---|
| **Breakfast** | 632.2 +/- 157.9 | 624.8 +/- 72.8 | 22.9 +/- 4.7 | 77.0 +/- 4.6 | 26.9 +/- 6.4 |
| **Lunch** | 520.1 +/- 72.3 | 554.0 +/- 31.4 | 16.0 +/- 3.2 | 82.2 +/- 9.7 | 22.7 +/- 4.8 |
| **Dinner** | 715.2 +/- 50.8 | 736.7 +/- 49.2 | 19.5 +/- 2.1 | 106.7 +/- 10.2 | 38.0 +/- 5.0 |
| **Snack** | 506.1 +/- 72.4 | 579.5 +/- 37.5 | 19.3 +/- 4.1 | 79.5 +/- 8.3 | 24.0 +/- 1.5 |
| **Beverage Only** | 197.5 +/- 93.7 | 2.6 +/- 1.3 | 0 +/- 0 | 0.657 +/- 0.3 | 0 +/- 0 |

C. Nutritional Information for Each Subject

| Subject | Date of Intake | Meal Name | Total Grams | Energy (kcal) | Total Fat (g) | Total Carbohydrate (g) | Total Protein (g) |
|---|---|---|---|---|---|---|---|
| 1 | 11/5/2015 | Dinner/Supper | 999.0 | 946.1 | 26.9 | 140.1 | 45.2 |
| 1 | 11/6/2015 | Breakfast | 879.4 | 716.4 | 23.7 | 95.1 | 30.3 |
| 1 | 11/6/2015 | Lunch | 509.2 | 691.5 | 18.2 | 108.5 | 33.1 |
| 1 | 11/6/2015 | Dinner/Supper | 921.2 | 995.5 | 32.4 | 128.2 | 51.7 |
| 1 | 11/7/2015 | Breakfast | 691.3 | 739.7 | 26.7 | 101.0 | 26.5 |
| 1 | 11/7/2015 | Lunch | 544.2 | 688.4 | 23.7 | 100.2 | 19.7 |
| 1 | 11/7/2015 | Dinner/Supper | 763.6 | 910.6 | 22.8 | 142.9 | 39.6 |
| 1 | 11/19/2015 | Dinner/Supper | 1011.4 | 957.1 | 27.1 | 138.3 | 45.3 |
| 1 | 11/19/2015 | Snack | 608.8 | 712.7 | 23.5 | 102.6 | 26.7 |
| 1 | 11/20/2015 | Beverage Only | 241.2 | 3.2 | 0.0 | 0.8 | 0.0 |
| 1 | 11/20/2015 | Lunch | 544.6 | 737.6 | 19.0 | 116.5 | 35.0 |
| 1 | 11/20/2015 | Dinner/Supper | 919.1 | 1002.4 | 31.9 | 130.7 | 52.1 |
| 1 | 11/20/2015 | Snack | 792.1 | 797.1 | 25.8 | 108.0 | 32.9 |
| 1 | 11/21/2015 | Beverage Only | 240.9 | 3.2 | 0.0 | 0.8 | 0.0 |
| 1 | 11/21/2015 | Lunch | 773.3 | 684.4 | 23.4 | 100.0 | 19.7 |
| 1 | 11/21/2015 | Dinner/Supper | 760.5 | 910.6 | 22.8 | 142.9 | 39.6 |
| 1 | 11/21/2015 | Snack | 445.9 | 724.4 | 26.9 | 96.3 | 26.7 |
| 2 | 1/6/2016 | Dinner/Supper | 1072.6 | 989.2 | 28.3 | 146.1 | 47.1 |
| 2 | 1/6/2016 | Snack | 499.3 | 803.3 | 26.5 | 116.7 | 29.0 |
| 2 | 1/7/2016 | Beverage Only | 220.7 | 2.8 | 0.0 | 0.7 | 0.0 |
| 2 | 1/7/2016 | Lunch | 509.6 | 684.0 | 17.9 | 107.6 | 32.4 |
| 2 | 1/7/2016 | Dinner/Supper | 922.4 | 1011.5 | 32.7 | 131.1 | 52.2 |
| 2 | 1/7/2016 | Snack | 714.1 | 805.4 | 25.4 | 110.4 | 33.3 |
| 2 | 1/8/2016 | Beverage Only | 241.0 | 3.2 | 0.0 | 0.8 | 0.0 |
| 2 | 1/8/2016 | Lunch | 762.3 | 681.4 | 23.4 | 99.3 | 19.6 |
| 2 | 1/8/2016 | Dinner/Supper | 721.1 | 839.7 | 19.1 | 136.0 | 36.3 |
| 2 | 1/8/2016 | Snack | 757.0 | 773.4 | 27.8 | 105.5 | 27.7 |
| 2 | 1/12/2016 | Dinner/Supper | 972.2 | 923.3 | 25.9 | 140.5 | 41.5 |
| 2 | 1/13/2016 | Breakfast | 943.0 | 783.9 | 24.3 | 107.4 | 33.2 |
| 2 | 1/13/2016 | Lunch | 523.3 | 711.6 | 18.3 | 111.9 | 34.3 |
| 2 | 1/13/2016 | Dinner/Supper | 912.7 | 970.6 | 29.9 | 130.2 | 49.0 |
| 2 | 1/14/2016 | Breakfast | 748.7 | 761.8 | 26.5 | 106.0 | 27.4 |
| 2 | 1/14/2016 | Lunch | 765.6 | 681.1 | 23.4 | 99.2 | 19.6 |
| 2 | 1/14/2016 | Dinner/Supper | 757.1 | 877.4 | 19.7 | 141.4 | 39.3 |

| Subject | Date of Intake | Meal Name | Total Grams | Energy (kcal) | Total Fat (g) | Total Carbohydrate (g) | Total Protein (g) |
|---|---|---|---|---|---|---|---|
| 3 | 3/18/2016 | Dinner/Supper | 975.4 | 925.9 | 26.4 | 143.2 | 38.5 |
| 3 | 3/18/2016 | Snack | 607.3 | 744.3 | 23.8 | 108.6 | 27.7 |
| 3 | 3/19/2016 | Beverage Only | 236.8 | 3.2 | 0.0 | 0.8 | 0.0 |
| 3 | 3/19/2016 | Lunch | 524.9 | 719.6 | 18.5 | 113.6 | 34.3 |
| 3 | 3/19/2016 | Dinner/Supper | 849.7 | 926.9 | 29.9 | 124.9 | 43.3 |
| 3 | 3/19/2016 | Snack | 520.1 | 594.8 | 19.5 | 82.1 | 23.0 |
| 3 | 3/20/2016 | Beverage Only | 237.2 | 3.2 | 0.0 | 0.8 | 0.0 |
| 3 | 3/20/2016 | Lunch | 725.7 | 619.7 | 20.5 | 93.3 | 16.8 |
| 3 | 3/20/2016 | Dinner/Supper | 671.2 | 828.8 | 21.3 | 132.9 | 32.0 |
| 3 | 3/20/2016 | Snack | 531.1 | 760.4 | 26.9 | 105.8 | 26.4 |
| 3 | 4/1/2016 | Dinner/Supper | 990.0 | 952.1 | 27.0 | 181.2 | 45.3 |
| 3 | 4/2/2016 | Breakfast | 932.6 | 745.0 | 22.4 | 106.9 | 28.7 |
| 3 | 4/2/2016 | Lunch | 532.3 | 719.6 | 18.4 | 113.6 | 34.6 |
| 3 | 4/2/2016 | Dinner/Supper | 916.2 | 994.9 | 31.8 | 129.2 | 51.9 |
| 3 | 4/3/2016 | Breakfast | 749.8 | 753.4 | 26.3 | 105.7 | 26.0 |
| 3 | 4/3/2016 | Lunch | 774.9 | 684.3 | 23.4 | 100.0 | 19.7 |
| 3 | 4/3/2016 | Dinner/Supper | 774.9 | 914.5 | 22.9 | 143.5 | 39.6 |
| 4 | 7/9/2016 | Dinner/Supper | 707.1 | 734.4 | 21.1 | 110.5 | 32.6 |
| 4 | 7/10/2016 | Breakfast | 761.9 | 579.9 | 18.3 | 80.0 | 23.5 |
| 4 | 7/10/2016 | Lunch | 465.3 | 562.5 | 13.1 | 88.7 | 26.7 |
| 4 | 7/10/2016 | Dinner/Supper | 729.9 | 724.6 | 20.0 | 96.1 | 41.7 |
| 4 | 7/11/2016 | Breakfast | 674.8 | 586.3 | 22.1 | 74.0 | 23.2 |
| 4 | 7/11/2016 | Lunch | 600.5 | 563.0 | 21.1 | 76.6 | 19.6 |
| 4 | 7/11/2016 | Dinner/Supper | 764.1 | 844.2 | 20.8 | 130.7 | 39.5 |
| 4 | 7/21/2016 | Dinner/Supper | 688.1 | 710.2 | 19.7 | 107.7 | 32.0 |
| 4 | 7/21/2016 | Snack | 564.2 | 567.9 | 18.6 | 84.1 | 22.4 |
| 4 | 7/22/2016 | Beverage Only | 233.2 | 3.2 | 0.0 | 0.8 | 0.0 |
| 4 | 7/22/2016 | Lunch | 469.4 | 583.7 | 13.3 | 92.2 | 28.1 |
| 4 | 7/22/2016 | Dinner/Supper | 734.2 | 757.0 | 22.1 | 98.0 | 43.6 |
| 4 | 7/22/2016 | Snack | 496.8 | 625.0 | 20.7 | 82.4 | 26.8 |
| 4 | 7/23/2016 | Beverage Only | 236.0 | 3.2 | 0.0 | 0.8 | 0.0 |
| 4 | 7/23/2016 | Lunch | 553.0 | 489.7 | 17.7 | 68.7 | 16.5 |
| 4 | 7/23/2016 | Dinner/Supper | 629.1 | 683.1 | 15.5 | 105.7 | 35.6 |
| 4 | 7/23/2016 | Snack | 450.5 | 603.5 | 22.5 | 76.5 | 24.4 |

| Subject | Date of Intake | Meal Name | Total Grams | Energy (kcal) | Total Fat (g) | Total Carbohydrate (g) | Total Protein (g) |
|---|---|---|---|---|---|---|---|
| 5 | 6/21/2016 | Dinner/Supper | 632.6 | 673.4 | 18.9 | 102.0 | 30.4 |
| 5 | 6/21/2016 | Breakfast | 761.1 | 767.2 | 32.2 | 78.1 | 39.8 |
| 5 | 6/22/2016 | Lunch | 411.0 | 561.9 | 13.0 | 88.7 | 27.0 |
| 5 | 6/22/2016 | Dinner/Supper | 767.2 | 777.4 | 21.8 | 103.0 | 44.4 |
| 5 | 6/23/2016 | Breakfast | 728.3 | 613.5 | 21.7 | 79.9 | 25.0 |
| 5 | 6/23/2016 | Lunch | 559.8 | 501.5 | 19.1 | 68.3 | 17.8 |
| 5 | 6/23/2016 | Dinner/Supper | 711.0 | 777.5 | 16.9 | 125.2 | 36.3 |
| 5 | 6/28/2016 | Dinner/Supper | 690.3 | 721.8 | 20.5 | 110.1 | 31.0 |
| 5 | 6/28/2016 | Snack | 561.1 | 523.7 | 17.7 | 74.6 | 22.4 |
| 5 | 6/29/2016 | Beverage Only | 236.0 | 3.2 | 0.0 | 0.8 | 0.0 |
| 5 | 6/29/2016 | Lunch | 424.4 | 575.1 | 13.3 | 91.2 | 26.8 |
| 5 | 6/29/2016 | Dinner/Supper | 703.4 | 692.8 | 19.0 | 95.3 | 36.7 |
| 5 | 6/29/2016 | Snack | 455.5 | 549.7 | 17.0 | 76.6 | 22.3 |
| 5 | 6/30/2016 | Beverage Only | 236.1 | 3.2 | 0.0 | 0.8 | 0.0 |
| 5 | 6/30/2016 | Lunch | 593.3 | 548.6 | 15.9 | 75.4 | 16.1 |
| 5 | 6/30/2016 | Dinner/Supper | 717.2 | 733.6 | 17.5 | 111.8 | 38.1 |
| 5 | 6/30/2016 | Snack | 421.5 | 632.4 | 24.1 | 78.9 | 25.2 |
| 6 | 2/3/2017 | Dinner/Supper | 625.6 | 711.4 | 20.7 | 101.1 | 36.1 |
| 6 | 2/3/2017 | Snack | 644.2 | 550.8 | 10.4 | 98.5 | 23.5 |
| 6 | 2/4/2017 | Beverage Only | 6.3 | 0.0 | 0.0 | 0.0 | 0.0 |
| 6 | 2/4/2017 | Lunch | 483.2 | 592.0 | 13.5 | 94.1 | 28.1 |
| 6 | 2/4/2017 | Dinner/Supper | 758.3 | 768.0 | 22.1 | 98.5 | 45.7 |
| 6 | 2/4/2017 | Snack | 514.8 | 561.2 | 20.7 | 69.0 | 24.7 |
| 6 | 2/5/2017 | Beverage Only | 237.9 | 3.2 | 0.0 | 0.8 | 0.0 |
| 6 | 2/5/2017 | Lunch | 596.9 | 553.6 | 20.7 | 75.4 | 19.6 |
| 6 | 2/5/2017 | Dinner/Supper | 751.6 | 776.4 | 18.3 | 119.7 | 38.9 |
| 6 | 2/5/2017 | Snack | 446.7 | 602.1 | 22.7 | 75.2 | 24.4 |
| 6 | 2/17/2017 | Dinner/Supper | 719.5 | 652.9 | 18.3 | 93.2 | 35.2 |
| 6 | 2/18/2017 | Breakfast | 461.0 | 628.4 | 21.3 | 81.0 | 26.9 |
| 6 | 2/18/2017 | Lunch | 478.0 | 580.3 | 13.4 | 92.4 | 26.8 |
| 6 | 2/18/2017 | Dinner/Supper | 820.6 | 809.2 | 23.1 | 104.4 | 48.0 |
| 6 | 2/19/2017 | Breakfast | 406.6 | 573.6 | 22.5 | 69.0 | 23.3 |
| 6 | 2/19/2017 | Lunch | 606.4 | 536.7 | 18.7 | 75.4 | 19.6 |
| 6 | 2/19/2017 | Dinner/Supper | 725.0 | 713.4 | 15.9 | 109.6 | 38.3 |

**Supplementary Table 2. Inclusion/Exclusion Criteria**

**Inclusion/Exclusion criteria**

<u>Inclusion Criteria --subject must</u>

- Be able to understand the study, provide written informed consent (in English), and be able to fill out the questionnaire
- Be male or female older than 18 years of age
- Have a normal BMI (20-25) or be obese (BMI more than 30)
- Have a normal basal glucose level (70-100 mg/dL)
- If female of childbearing potential, have a negative pregnancy test on study day

<u>Exclusion Criteria-- subject must not</u>

- Be pregnant or lactating
- Have known sleep, metabolic (e.g., diabetes), or gastro-intestinal disorders except obesity
- Had alcohol less than 24 hours before admission
- Require assistance with activities of daily living
- Have difficulty swallowing
- Be unable to complete a food and sleep diary
- Be smokers

**Supplementary Table 3. Subject Recruitment Questionnaire**

1. When do you sleep on a typical day?

   a) Bedtime:________________(e.g., 10:00-11:00 PM)

   b) Wake-up time____________(e.g., 06:00-7:00 AM)

2. The following questions concern your typical meal times and food intake:

   a) Do you normally eat breakfast? ___Yes ___No

   If you answered Yes, what time do you usually eat breakfast? ___________

   Please give an example of what you might eat for a typical breakfast (e.g., piece of toast, bowl of cereal, or bacon, eggs, & toast, etc).

   ___________________________________________________

   b) Do you normally eat lunch? ___Yes ___No

   If you answered Yes, what time do you usually eat lunch? ______

   Please give an example of what you might eat for a typical lunch

   ___________________________________________________

   c) Do you normally eat dinner? ___Yes ___No

   If you answered Yes, what time do you usually eat dinner? ______

   Is dinner generally your largest meal of the day? ___Yes ___No

   d) Do you frequently eat snacks between meals or after dinner? ___Yes ___No

   If you answered Yes, what time(s) do you usually have your snacks relative to:

   your main meals (check all that apply) ?

   ______In the morning between breakfast and lunch

   ______In the afternoon between lunch and dinner

   ______After dinner

   e) How many caffeinated drinks do you drink daily? (Please answer in terms of the number of cups of coffee or tea or number of portions of caffeinated soft drinks per day): ________________________________________________

   f) What is your average alcohol intake? (Please answer in terms of the number of alcohol-containing drinks per week): ____________________________________

g) Do you have any dietary restrictions? ___Yes ___No

If you answered Yes, please describe what they are. ______________________

______________________________________________________________

3. Do you routinely exercise ___Yes ___No

If you answered Yes, please answer the following questions:

a) What type(s) of exercise do you do (e.g., jogging, swimming, yoga, etc.)

____________________________________________________

b) When (relative to your mealtimes) do you normally exercise?

____________________________

c) In a typical week, how many days do you exercise?

____1-2 ____3-4 ____5-7

4. Have you ever been diagnosed with any of the following diseases: ___Yes ___No

Esophageal stricture

Diverticulosis

Inflammatory bowel disease (IBD),

Peptic ulcer disease

Crohn's disease

Ulcerative colitis

5. Have you ever had gastrointestinal surgery: ___Yes ___No

6. The protocol for this study involves swallowing a capsule the size of a large vitamin. Do you have any difficulty swallowing: ___Yes ___No

7. Do you have any chronic medical problems not listed above (e.g., diabetes, high blood pressure, asthma, etc)

___Yes ___No

If you answered yes, please describe them._______________________

8. Are you currently taking any medications? ___Yes ___No

If you answered yes, please describe them. ________________________

____________________________________________________________________

9. Do you have any sleep disorders (e.g., sleep apnea, insomnia, sleep walking, restless leg syndrome, etc.)?

___Yes ___No

If you answered yes, please describe them. ________________________

____________________________________________________________________

10. Please complete the following demographic information:

Gender: _____male _____female

Age: _____ years

Ethnic group: _____ African-American

_____ Asian

_____ Caucasian

_____ Hispanic

_____ Other

**Supplementary Table 4. Hour by Hour Mixed Model Analysis**

**A. RER**

| RER Pairwise | Hour | Estimate | SE | df | t.ratio | p.value |
|---|---|---|---|---|---|---|
| Breakfast - Snack | 15 | -0.0219 | 0.0131 | 5 | 1.6723 | 0.1553 |
| Breakfast - Snack | 16 | -0.0191 | 0.0129 | 5 | 1.4761 | 0.1999 |
| Breakfast - Snack | 17 | 0.0099 | 0.0118 | 5 | -0.8392 | 0.4396 |
| Breakfast - Snack | 18 | 0.0023 | 0.0117 | 5 | -0.1951 | 0.853 |
| Breakfast - Snack | 19 | -0.0171 | 0.0117 | 5 | 1.4586 | 0.2045 |
| Breakfast - Snack | 20 | -0.0125 | 0.0117 | 5 | 1.065 | 0.3356 |
| Breakfast - Snack | 21 | -0.0083 | 0.0117 | 5 | 0.7098 | 0.5095 |
| Breakfast - Snack | 22 | -0.0522 | 0.0116 | 5 | 4.4943 | 0.0064 |
| Breakfast - Snack | 23 | -0.0536 | 0.0116 | 5 | 4.6134 | 0.0058 |
| Breakfast - Snack | 0 | -0.054 | 0.0116 | 5 | 4.6455 | 0.0056 |
| Breakfast - Snack | 1 | -0.0503 | 0.0117 | 5 | 4.2948 | 0.0078 |
| Breakfast - Snack | 2 | -0.0515 | 0.0117 | 5 | 4.3987 | 0.007 |
| Breakfast - Snack | 3 | -0.0308 | 0.0117 | 5 | 2.6279 | 0.0466 |
| Breakfast - Snack | 4 | -0.0265 | 0.0117 | 5 | 2.2608 | 0.0733 |
| Breakfast - Snack | 5 | -0.0308 | 0.0117 | 5 | 2.6299 | 0.0465 |
| Breakfast - Snack | 6 | -0.0199 | 0.0118 | 5 | 1.6892 | 0.152 |
| Breakfast - Snack | 7 | -0.0069 | 0.0125 | 5 | 0.5527 | 0.6043 |
| Breakfast - Snack | 8 | 0.0299 | 0.0131 | 5 | -2.2847 | 0.0711 |
| Breakfast - Snack | 9 | 0.032 | 0.0131 | 5 | -2.4491 | 0.058 |
| Breakfast - Snack | 10 | 0.0138 | 0.0133 | 5 | -1.0425 | 0.3449 |
| Breakfast - Snack | 11 | 0.0157 | 0.0131 | 5 | -1.2025 | 0.283 |
| Breakfast - Snack | 12 | 0.0198 | 0.0131 | 5 | -1.5133 | 0.1906 |
| Breakfast - Snack | 13 | -0.0083 | 0.0131 | 5 | 0.632 | 0.5551 |
| Breakfast - Snack | 14 | -0.0134 | 0.0131 | 5 | 1.0277 | 0.3512 |

**B. Activity**

| Activity Pairwise | Hour | Estimate | SE | df | t.ratio | p.value |
|---|---|---|---|---|---|---|
| Breakfast - Snack | 15 | 1.2773 | 0.4966 | 4 | -2.5722 | 0.0618 |
| Breakfast - Snack | 16 | -0.7947 | 0.4966 | 4 | 1.6003 | 0.1848 |
| Breakfast - Snack | 17 | -0.3094 | 0.4066 | 4 | 0.7609 | 0.4891 |
| Breakfast - Snack | 18 | 0.4559 | 0.4066 | 4 | -1.1211 | 0.325 |
| Breakfast - Snack | 19 | -1.0109 | 0.4066 | 4 | 2.4861 | 0.0678 |
| Breakfast - Snack | 20 | -0.449 | 0.4066 | 4 | 1.1043 | 0.3314 |
| Breakfast - Snack | 21 | -0.3579 | 0.4066 | 4 | 0.8801 | 0.4285 |
| Breakfast - Snack | 22 | -1.3114 | 0.4066 | 4 | 3.2253 | 0.0321 |
| Breakfast - Snack | 23 | -0.164 | 0.4066 | 4 | 0.4033 | 0.7074 |
| Breakfast - Snack | 0 | -0.3138 | 0.4066 | 4 | 0.7717 | 0.4834 |
| Breakfast - Snack | 1 | 0.36 | 0.4066 | 4 | -0.8854 | 0.426 |

| | | | | | | |
|---|---|---|---|---|---|---|
| Breakfast - Snack | 2 | -0.488 | 0.4066 | 4 | 1.2002 | 0.2963 |
| Breakfast - Snack | 3 | -0.0864 | 0.4066 | 4 | 0.2126 | 0.842 |
| Breakfast - Snack | 4 | -0.1766 | 0.4066 | 4 | 0.4343 | 0.6865 |
| Breakfast - Snack | 5 | -0.1878 | 0.4066 | 4 | 0.462 | 0.6681 |
| Breakfast - Snack | 6 | -0.2409 | 0.4066 | 4 | 0.5925 | 0.5854 |
| Breakfast - Snack | 7 | 0.4582 | 0.4066 | 4 | -1.1269 | 0.3228 |
| Breakfast - Snack | 8 | 0.7877 | 0.4966 | 4 | -1.5863 | 0.1879 |
| Breakfast - Snack | 9 | -0.5758 | 0.4966 | 4 | 1.1596 | 0.3107 |
| Breakfast - Snack | 10 | 0.298 | 0.5102 | 4 | -0.5841 | 0.5905 |
| Breakfast - Snack | 11 | 0.0051 | 0.4966 | 4 | -0.0102 | 0.9923 |
| Breakfast - Snack | 12 | -0.4765 | 0.4966 | 4 | 0.9595 | 0.3917 |
| Breakfast - Snack | 13 | 0.7597 | 0.4966 | 4 | -1.53 | 0.2008 |
| Breakfast - Snack | 14 | 0.6023 | 0.4966 | 4 | -1.2128 | 0.2919 |

**C. Temperature**

| Temperature Pairwise | Hour | Estimate | SE | df | t.ratio | p.value |
|---|---|---|---|---|---|---|
| Breakfast - Snack | 15 | 0.0525 | 0.143 | 5 | -0.367 | 0.7286 |
| Breakfast - Snack | 16 | 0.5505 | 0.1357 | 5 | -4.0568 | 0.0098 |
| Breakfast - Snack | 17 | -0.1352 | 0.1197 | 5 | 1.1297 | 0.3099 |
| Breakfast - Snack | 18 | -0.055 | 0.114 | 5 | 0.4824 | 0.6499 |
| Breakfast - Snack | 19 | -0.0562 | 0.1106 | 5 | 0.5084 | 0.6328 |
| Breakfast - Snack | 20 | -0.106 | 0.1106 | 5 | 0.9587 | 0.3817 |
| Breakfast - Snack | 21 | -0.081 | 0.1106 | 5 | 0.7331 | 0.4964 |
| Breakfast - Snack | 22 | -0.0436 | 0.1106 | 5 | 0.3939 | 0.7099 |
| Breakfast - Snack | 23 | -0.073 | 0.1121 | 5 | 0.6511 | 0.5437 |
| Breakfast - Snack | 0 | -0.1591 | 0.1106 | 5 | 1.4391 | 0.2096 |
| Breakfast - Snack | 1 | -0.1283 | 0.1121 | 5 | 1.1442 | 0.3043 |
| Breakfast - Snack | 2 | -0.0812 | 0.1106 | 5 | 0.7342 | 0.4958 |
| Breakfast - Snack | 3 | -0.0887 | 0.1121 | 5 | 0.7912 | 0.4647 |
| Breakfast - Snack | 4 | -0.1382 | 0.1139 | 5 | 1.2139 | 0.279 |
| Breakfast - Snack | 5 | -0.0937 | 0.1139 | 5 | 0.8225 | 0.4482 |
| Breakfast - Snack | 6 | -0.0243 | 0.1139 | 5 | 0.213 | 0.8398 |
| Breakfast - Snack | 7 | -0.0651 | 0.1197 | 5 | 0.5442 | 0.6097 |
| Breakfast - Snack | 8 | 0.002 | 0.1365 | 5 | -0.0148 | 0.9888 |
| Breakfast - Snack | 9 | 0.0344 | 0.1401 | 5 | -0.2459 | 0.8155 |
| Breakfast - Snack | 10 | 0.092 | 0.1553 | 5 | -0.5926 | 0.5792 |
| Breakfast - Snack | 11 | -0.025 | 0.1592 | 5 | 0.1569 | 0.8814 |
| Breakfast - Snack | 12 | 0.0777 | 0.143 | 5 | -0.5435 | 0.6102 |
| Breakfast - Snack | 13 | 0.0547 | 0.143 | 5 | -0.3823 | 0.7179 |
| Breakfast - Snack | 14 | 0.036 | 0.143 | 5 | -0.2516 | 0.8113 |

**D. Metabolic Rate**

| Metabolic Rate Pairwise | Hour | Estimate | SE | df | t.ratio | p.value |
|---|---|---|---|---|---|---|
| Breakfast - Snack | 15 | 0.068 | 0.0771 | 5 | -0.8814 | 0.4185 |
| Breakfast - Snack | 16 | 0.0887 | 0.0759 | 5 | -1.1692 | 0.295 |
| Breakfast - Snack | 17 | 0.1747 | 0.0667 | 5 | -2.617 | 0.0473 |
| Breakfast - Snack | 18 | 0.1899 | 0.066 | 5 | -2.8778 | 0.0347 |
| Breakfast - Snack | 19 | 0.031 | 0.066 | 5 | -0.47 | 0.6582 |
| Breakfast - Snack | 20 | 0.1421 | 0.066 | 5 | -2.1528 | 0.0839 |
| Breakfast - Snack | 21 | 0.0552 | 0.066 | 5 | -0.836 | 0.4413 |
| Breakfast - Snack | 22 | -0.3364 | 0.0652 | 5 | 5.1592 | 0.0036 |
| Breakfast - Snack | 23 | -0.1839 | 0.0652 | 5 | 2.8196 | 0.0371 |
| Breakfast - Snack | 0 | -0.0874 | 0.0652 | 5 | 1.3409 | 0.2377 |
| Breakfast - Snack | 1 | -0.0669 | 0.066 | 5 | 1.0134 | 0.3574 |
| Breakfast - Snack | 2 | -0.0592 | 0.066 | 5 | 0.8974 | 0.4107 |
| Breakfast - Snack | 3 | 0.0151 | 0.066 | 5 | -0.2283 | 0.8285 |
| Breakfast - Snack | 4 | 0.0468 | 0.066 | 5 | -0.709 | 0.51 |
| Breakfast - Snack | 5 | 0.0377 | 0.066 | 5 | -0.5706 | 0.593 |
| Breakfast - Snack | 6 | 0.052 | 0.0668 | 5 | -0.7787 | 0.4714 |
| Breakfast - Snack | 7 | 0.2516 | 0.0748 | 5 | -3.3623 | 0.0201 |
| Breakfast - Snack | 8 | 0.218 | 0.0771 | 5 | -2.8262 | 0.0368 |
| Breakfast - Snack | 9 | 0.2154 | 0.0771 | 5 | -2.792 | 0.0384 |
| Breakfast - Snack | 10 | 0.1684 | 0.0787 | 5 | -2.1405 | 0.0853 |
| Breakfast - Snack | 11 | 0.1639 | 0.0771 | 5 | -2.1246 | 0.087 |
| Breakfast - Snack | 12 | 0.0911 | 0.0771 | 5 | -1.1807 | 0.2908 |
| Breakfast - Snack | 13 | 0.143 | 0.0771 | 5 | -1.8543 | 0.1229 |
| Breakfast - Snack | 14 | 0.2571 | 0.0771 | 5 | -3.3332 | 0.0207 |

**E. Carbohydrate Oxidation**

| Carbohydrate Oxidation Pairwise | hour | estimate | SE | df | t.ratio | p.value |
|---|---|---|---|---|---|---|
| Breakfast - Snack | 15 | -0.0258 | 0.0225 | 5 | 1.1495 | 0.3023 |
| Breakfast - Snack | 16 | -0.0207 | 0.0221 | 5 | 0.9377 | 0.3915 |
| Breakfast - Snack | 17 | 0.0309 | 0.0197 | 5 | -1.5704 | 0.1771 |
| Breakfast - Snack | 18 | 0.0309 | 0.0195 | 5 | -1.5817 | 0.1746 |
| Breakfast - Snack | 19 | -0.024 | 0.0195 | 5 | 1.2298 | 0.2735 |
| Breakfast - Snack | 20 | -0.0018 | 0.0195 | 5 | 0.0947 | 0.9282 |
| Breakfast - Snack | 21 | -0.0082 | 0.0195 | 5 | 0.4182 | 0.6931 |
| Breakfast - Snack | 22 | -0.1342 | 0.0193 | 5 | 6.9532 | 9.00E-04 |
| Breakfast - Snack | 23 | -0.0978 | 0.0193 | 5 | 5.0662 | 0.0039 |
| Breakfast - Snack | 0 | -0.0764 | 0.0193 | 5 | 3.9563 | 0.0108 |
| Breakfast - Snack | 1 | -0.0721 | 0.0195 | 5 | 3.6973 | 0.014 |
| Breakfast - Snack | 2 | -0.0732 | 0.0195 | 5 | 3.752 | 0.0133 |

| | | | | | | |
|---|---|---|---|---|---|---|
| Breakfast - Snack | 3 | -0.0398 | 0.0195 | 5 | 2.0397 | 0.0969 |
| Breakfast - Snack | 4 | -0.0282 | 0.0195 | 5 | 1.4434 | 0.2085 |
| Breakfast - Snack | 5 | -0.0367 | 0.0195 | 5 | 1.8788 | 0.1191 |
| Breakfast - Snack | 6 | -0.0245 | 0.0197 | 5 | 1.2428 | 0.2691 |
| Breakfast - Snack | 7 | 0.0115 | 0.0218 | 5 | -0.5267 | 0.6209 |
| Breakfast - Snack | 8 | 0.0646 | 0.0225 | 5 | -2.8769 | 0.0347 |
| Breakfast - Snack | 9 | 0.0698 | 0.0225 | 5 | -3.1083 | 0.0266 |
| Breakfast - Snack | 10 | 0.0504 | 0.0229 | 5 | -2.2047 | 0.0786 |
| Breakfast - Snack | 11 | 0.0468 | 0.0225 | 5 | -2.0845 | 0.0915 |
| Breakfast - Snack | 12 | 0.0476 | 0.0225 | 5 | -2.1187 | 0.0877 |
| Breakfast - Snack | 13 | 0.0117 | 0.0225 | 5 | -0.523 | 0.6233 |
| Breakfast - Snack | 14 | 0.0148 | 0.0225 | 5 | -0.6571 | 0.5402 |

**F. Lipid Oxidation**

| Lipid Oxidation Pairwise | Hour | Estimate | SE | df | t.ratio | p.value |
|---|---|---|---|---|---|---|
| Breakfast - Snack | 15 | 0.0172 | 0.0079 | 5 | -2.1771 | 0.0814 |
| Breakfast - Snack | 16 | 0.0174 | 0.0078 | 5 | -2.2315 | 0.076 |
| Breakfast - Snack | 17 | 0 | 0.007 | 5 | -7.00E-04 | 0.9994 |
| Breakfast - Snack | 18 | 0.0076 | 0.0069 | 5 | -1.1047 | 0.3196 |
| Breakfast - Snack | 19 | 0.0127 | 0.0069 | 5 | -1.8343 | 0.1261 |
| Breakfast - Snack | 20 | 0.0155 | 0.0069 | 5 | -2.2461 | 0.0746 |
| Breakfast - Snack | 21 | 0.009 | 0.0069 | 5 | -1.2967 | 0.2513 |
| Breakfast - Snack | 22 | 0.0178 | 0.0068 | 5 | -2.6009 | 0.0482 |
| Breakfast - Snack | 23 | 0.0193 | 0.0068 | 5 | -2.8271 | 0.0368 |
| Breakfast - Snack | 0 | 0.021 | 0.0068 | 5 | -3.0619 | 0.028 |
| Breakfast - Snack | 1 | 0.0214 | 0.0069 | 5 | -3.0991 | 0.0269 |
| Breakfast - Snack | 2 | 0.0226 | 0.0069 | 5 | -3.2752 | 0.0221 |
| Breakfast - Snack | 3 | 0.0172 | 0.0069 | 5 | -2.4921 | 0.055 |
| Breakfast - Snack | 4 | 0.0159 | 0.0069 | 5 | -2.3075 | 0.0691 |
| Breakfast - Snack | 5 | 0.0183 | 0.0069 | 5 | -2.6536 | 0.0452 |
| Breakfast - Snack | 6 | 0.0151 | 0.007 | 5 | -2.1623 | 0.0829 |
| Breakfast - Snack | 7 | 0.0218 | 0.0077 | 5 | -2.8242 | 0.0369 |
| Breakfast - Snack | 8 | -0.0027 | 0.0079 | 5 | 0.3448 | 0.7442 |
| Breakfast - Snack | 9 | -0.005 | 0.0079 | 5 | 0.6378 | 0.5517 |
| Breakfast - Snack | 10 | -0.0023 | 0.0081 | 5 | 0.2818 | 0.7894 |
| Breakfast - Snack | 11 | -0.0014 | 0.0079 | 5 | 0.1719 | 0.8703 |
| Breakfast - Snack | 12 | -0.0092 | 0.0079 | 5 | 1.1675 | 0.2957 |
| Breakfast - Snack | 13 | 0.0103 | 0.0079 | 5 | -1.2969 | 0.2513 |
| Breakfast - Snack | 14 | 0.021 | 0.0079 | 5 | -2.6474 | 0.0456 |
